# Metabolic plasticity of serine metabolism is crucial for cGAS/STING-signalling and innate immune response to viral infections in the gut

**DOI:** 10.1101/2022.05.17.492340

**Authors:** Björn Becker, Felix Wottawa, Mohamed Bakr, Eric Koncina, Lisa Mayr, Julia Kugler, Guang Yang, Samuel J Windross, Laura Neises, Neha Mishra, Danielle Harris, Florian Tran, Lina Welz, Julian Schwärzler, Zoltán Bánki, Stephanie T Stengel, Go Ito, Christina Krötz, Olivia I Coleman, Christian Jaeger, Dirk Haller, Søren R Paludan, Richard Blumberg, Arthur Kaser, Luka Cicin-Sain, Stefan Schreiber, Timon E. Adolph, Elisabeth Letellier, Philip Rosenstiel, Johannes Meiser, Konrad Aden

## Abstract

Inflammatory bowel diseases (IBD) are characterized by chronic relapsing inflammation of the gastrointestinal tract. While the molecular causality between endoplasmic reticulum (ER) stress and intestinal inflammation is widely accepted, the metabolic consequences of chronic ER-stress on the pathophysiology of IBD remain unclear. By using *in vitro*, *ex vivo*, *in vivo* mouse models and patient datasets, we identified a distinct polarisation of the mitochondrial one-carbon (1C) metabolism and a fine-tuning of the amino acid uptake in intestinal epithelial cells tailored to support GSH and NADPH metabolism upon chronic ER-stress. This metabolic phenotype strongly correlates with IBD severity and therapy-response. Mechanistically, we uncover that both chronic ER-stress and serine limitation disrupt cGAS/STING-signalling, impairing the epithelial response against viral and bacterial infection, fuelling experimental enteritis. Consequently, antioxidant treatment restores STING function and virus control. Collectively, our data highlight the importance of the plasticity of serine metabolism to allow proper cGAS/STING-signalling and innate immune responses upon chronic inflammation in the gut.

## Introduction

IBD, classically divided into ulcerative colitis (UC) and Crohn’s disease (CD), describes a group of chronic disorders defined by relapsing and remitting episodes of gastrointestinal inflammation. Although the introduction of targeted inhibition of pro-inflammatory pathways (e.g. anti-TNF, anti-IL12/23) has significantly improved patient care, disease-associated complications such as IBD-related hospitalisations and surgery still contribute to the major socioeconomic burden of IBD (*1*). In the past years, IBD-related cytomegalovirus (CMV) infections have seen a steep rise and increase the risk of hospitalisation and proctocolectomy (2).

Several studies already demonstrated that the deregulation of ER-stress signalling pathways in intestinal epithelial cells (IECs) is linked to IBD initiation and progression (*2–6*). ER-stress describes a state of disrupted cellular homeostasis, which occurs when the balance between ER-folding capacity and protein load is disturbed (*7, 8*). Thereby, the consequences of ER-stress on intestinal homeostasis appear to be multifaceted, ranging from impaired mucosal barrier function over altered intestinal microbial composition to uncontrolled immune responses (*9–11*). Especially, IECs with a high secretory demand such as Paneth and Goblet cells are highly susceptible to changes in ER homeostasis (*5, 6*). Ultimately, ER-stress triggers a detrimental inflammatory reaction that drives mucosal tissue damage in human IBD patients (*12, 13*). Usually, to restore ER homeostasis, cells activate the unfolded protein response (UPR) which consists of three major pathways (specifically, the IRE1/XBP1-, PERK/ATF4- and ATF6-pathway) (*7*). Single-nucleotide-polymorphisms (SNPs) in *X-box binding protein 1* (*XBP1*) cause an ER-stress sensitive variant resulting in increased IBD susceptibility and intestinal tumorigenesis (*14, 15*).

In contrast to the pathophysiological consequences, the metabolic consequences to impaired ER homeostasis in the context of IBD remain largely unattended. Besides the clearance of misfolded proteins, ER-stressed cells have to reprogram their metabolism to handle the increased oxidative burden. For instance, cells can maximise the metabolic output towards Reduction/Oxidation (RedOx) control by utilising available carbon sources differently. In this regards, the serine and one-carbon (1C) metabolism are of great important to provide the building block glycine for the synthesis of the most abundant antioxidant glutathione (GSH) (*16*). Since GSH synthesis also requires the amino acids cysteine and glutamate, cells need to additionally fine-tune the availability of these amino acids (*17*). Importantly, a prolonged disequilibrium of the RedOx homeostasis is one of the notorious features of patients suffering from chronic inflammatory disease (*18, 19*).

Another important aspect is the association between impaired RedOx homeostasis and increased lipid peroxidation (LPO) (*20*). Recent studies indicate that by-products of LPO reactions (e.g. 4-HNE) interfere with the activation of the cGAS/STING pathway (*21*). In general, this pathway allows the initiation of the host defence against pathogens by directly sensing cytosolic double-stranded DNA (dsDNA) derived from viruses (e.g. CMV) or cyclic dinucleotides derived from intracellular bacteria (e.g. *Listeria monocytogenes*) (*22–24*). After binding of cytosolic dsDNA and subsequent cGAMP production by cGAS, STING is activated and transported from the ER through the ER-Golgi intermediate compartment (ERGIC) into the Golgi-apparatus. Here, STING recruits Tank-binding-kinase 1 (TBK1) and interferon-regulatory-factor 3 (IRF3), controlling type-I IFN (e.g. IFN-β) production and secretion. IFN-β then induces a variety of antiviral, immunomodulatory or anti-proliferative interferon-stimulated genes (ISGs) such as *CXCL10* (*24*). As such, cGAS/STING-signalling is critically involved in type-I IFN induction during cytomegalovirus (CMV) infection, and CMV actively antagonises STING-mediated IFN induction, highlighting the potential role of STING as a crucial first line of defence against CMV infection (*25–27*). Interestingly, meta-analyses of retrospective clinical data of IBD patients indicate that the disease extent and disease severity directly correlate with CMV colitis risk. However, the underlying molecular mechanisms that drive susceptibility to CMV infection in IBD patients remain enigmatic (*28*). Due to its localisation as an ER-resident protein, it is postulated that STING function is closely intertwined with ER homeostasis. However, to which extent disturbance of the UPR directly affects cGAS/STING-sensing is not known. Several studies support the notion that ER-stress is an important upstream modulator of STING activity in response to PAMPs and DAMPs (*29, 30*) and pro-regenerative IL-22 signalling (*31*).

In this study, we combined metabolic profiling with transcriptome analyses of IBD cohorts and discovered a distinct metabolic reprogramming towards GSH metabolism in the intestinal epithelium upon chronic ER-stress. This metabolic phenotype correlates with IBD severity and therapy-response status. Despite the metabolic rewiring towards RedOx control, ER-stressed IECs show an imbalanced RedOx homeostasis suppressing the initiation of cGAS/STING-signalling in response to invading viruses and bacteria. Using pharmacological interventions, we further underpinned the essential role of the serine metabolism regarding cGAS/STING signalling upon chronic ER-stress. Our findings therefore shed new lights on the underlying mechanisms driving increased susceptibility of IBD patients against CMV infections.

## Results

### Chronic ER-stress promotes glutathione synthesis by adapting amino acid uptake in IECs

Unresolved ER-stress represents a critical hallmark of chronic intestinal inflammation and contributes to the pathogenesis of IBD (*10, 32*). As such, defects of components of the UPR machinery are linked to IBD development (*5, 33, 34*). Since the loss of the IBD risk gene *XBP1*/*Xbp1* causes ER-stress and drives intestinal inflammation in mice and humans (*5*), we studied the metabolic consequences of Xbp1 deficiency in IECs (i*Xbp1*) by using different metabolic read-outs (Fig. 1A).

**Fig. 1:**
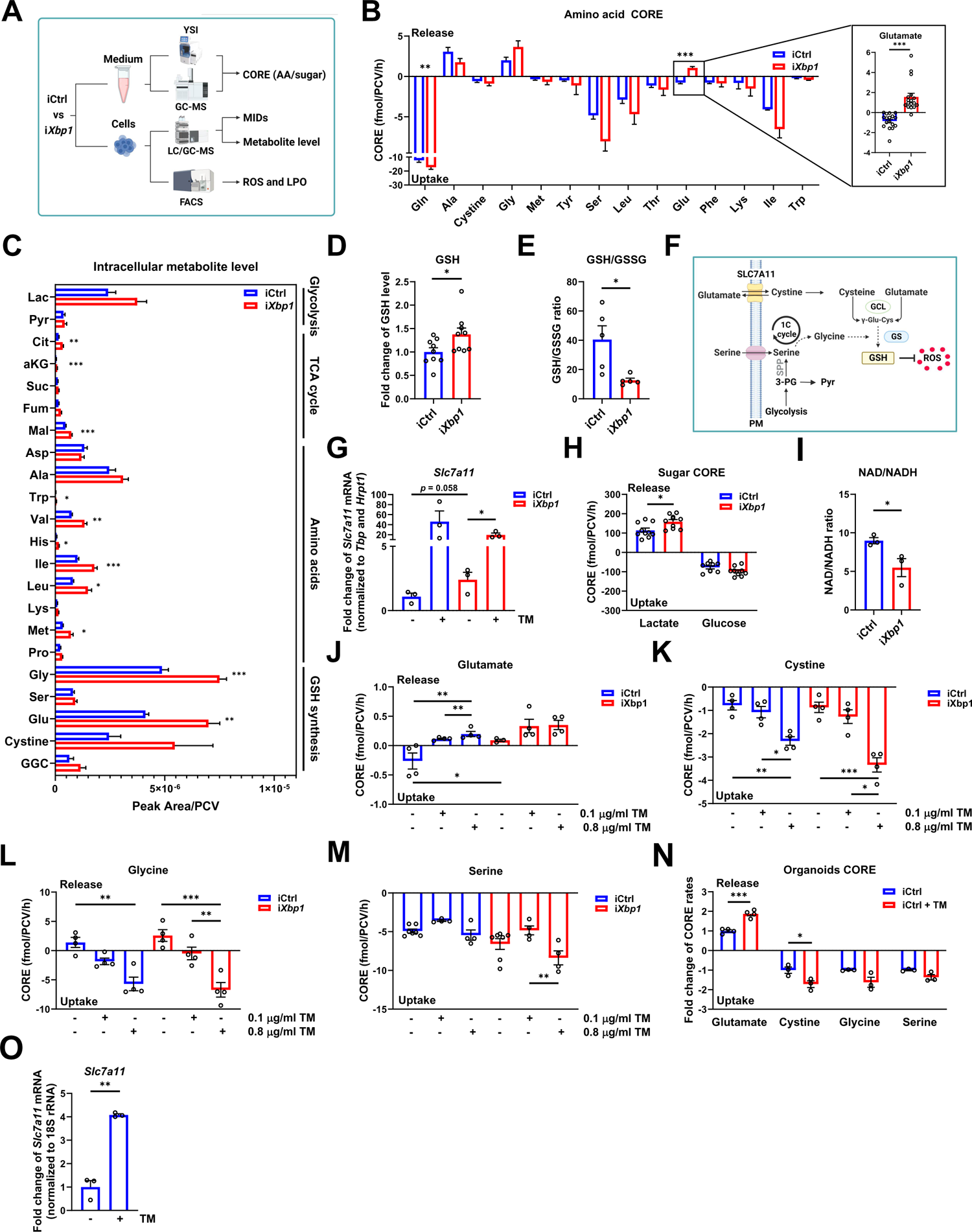
ER-stress induces a distinct ROS-dependent metabolic reprogramming of the amino acid metabolism to support GSH synthesis in IECs. **(A)** Schematic overview of the experimental workflow used to analyse metabolic and cellular changes in wild-type (iCtrl) and chronically ER-stressed (i*Xbp1*) Mode-K cells. **(B)** *In vitro* uptake rates of amino acids (left) and zoom-in of glutamate (right). Absolute consumption and release rates (CORE) in fmol/PCV/h are displayed (n ≥ 3). **(C)** Intracellular metabolite level (Peak Area/PCV) in iCtrl and i*Xbp1* cells (n ≥ 3). **(D)** Intracellular GSH level in iCtrl and i*Xbp1* cells (n=9, paired t-test). **(E)** GSH/GSSG ratios in iCtrl and i*Xbp1* cells (n=3, paired t-test). **(F)** Schematic illustration of the GSH synthesis pathway. (GCL: glutamate-cysteine ligase; GS: glutathione synthetase, SSP: serine *de novo* synthesis pathway; PM: plasma membrane). **(G)** *Slc7a11* mRNA expression in iCtrl and i*Xbp1* cells treated with (+) or without (-) TM (n=3). **(H)** Absolute CORE rates in fmol/PCV/h of glucose and lactate in iCtrl (n = 10) and i*Xbp1* cells (n=9) **(I)** NAD/NADH ratio in iCtrl and i*Xbp1* cells (n = 3). **(J-M)** Absolute CORE rates of (J) glutamate, (K) cysteine, (L) glycine and (M) serine in iCtrl and i*Xbp1* cells in response to TM treatment (n=4, Two-way ANOVA). **(N)** Absolute CORE rates of the indicated amino acids in murine SI organoids cultures (derived from *Xbp1*^fl/fl^ mice) in the presence or absence of 0.1 µg/ml TM. Representative experiment is shown. **(O)** *Slc7a11* mRNA expression in murine SI organoids (derived from *Xbp1*^fl/fl^ mice) treated with (+) or without (-) 0.1 µg/ml TM (n=3, paired t-test). \**P*<0.05, \*\**P*<0.01, \*\*\**P*<0.001.

While ^13^C flux analysis did not reveal striking differences in the central carbon metabolism of chronically ER-stressed IECs (Fig S1A-D), exchange rates of distinct amino acids were heavily altered. Xbp1 loss results in increased glutamine consumption and a striking switch from glutamate consumption towards glutamate release. Additionally, we also observed a general trend towards increased amino acid uptake under *Xbp1* deficiency (Fig. 1B). Intracellular metabolic steady-state levels of various metabolites including different amino acids and TCA cycle intermediates were elevated. Most consistently, i*Xbp1* cells showed a strong increase of metabolites related to glutathione (GSH) synthesis, such as glycine, glutamate and cystine (Fig. 1C). In line with this notion, intracellular GSH level were increased in i*Xbp1* cells (Fig. 1D). Since the primordial function of GSH is to counterbalance free ROS-derived radicals, the reduced GSH over GSSG ratio suggests an unbalanced RedOx homeostasis under *Xbp1*-deficiency (Fig. 1E). Additionally, cells can increase intracellular cysteine availability via the cystine/glutamate antiporter Slc7a11 to support GSH synthesis (*35*) (Fig. 1F). In line with this, we observed *Slc7a11* upregulation in response to ER-stress (Fig 1G). Moreover, i*Xbp1* cells released more lactate and showed a reduced NAD/NADH ratio compared to wild-type counterparts (iCtrl,) (Fig 1H, I). Overall, the results indicate a metabolic plasticity towards GSH metabolism to counterbalance reactive oxygen species (ROS).

To investigate whether the observed effects are generalizable to ER-stress, we used the ER-stress inducer Tunicamycin (TM) in iCtrl cells and were able to phenocopy the glutamate release phenotype of i*Xbp1* cells in a dose-dependent manner (Fig. 1J). In line with this finding, we also observed increased cystine uptake rates upon TM treatment. Surprisingly, TM treatment triggers a switch from glycine release towards glycine uptake (Fig. 1K+L). In contrast, serine consumption was not altered and only elevated at higher TM doses in i*Xbp1* cells (Fig. 1M). Generally, serine catabolism provides sufficient amounts of glycine resulting in net excretion of glycine (*36, 37*). However, in context of ER-stress, serine-derived glycine seems not to be sufficient for IECs resulting in a switch from net excretion to net consumption. Similar results were also observed in intestinal organoids (Fig. 1N+O). Interestingly, the obtained data suggest a higher baseline stress level in organoids compared to the 2D cell culture model.

Overall, our findings demonstrate that chronically ER-stressed IECs promote GSH synthesis by modulating the uptake of GSH-relevant amino acids from the extracellular space.

### Metabolic reprogramming of the 1C metabolism fuels the NADPH pool upon ER-stress

In addition to providing glycine for GSH synthesis, the 1C metabolism can directly supply NADPH thereby contributing to cellular RedOx balance (*38*). Depending on the cellular demands, cells can convert 1C units via MTHFD1L into formate generating ATP or fully oxidise 1C units via ALDH1L1/L2 to CO_2_ generating NADPH (Fig. 2A+S2A) (*16*). Since previous studies indicate that ER-stress increases intracellular ROS levels (*39, 40*), we postulated a metabolic rewiring towards ALDH1L1 or ALDH1L2 to enhance the reducing capacity in response to ER-stress. Increased NADPH levels and unchanged formate release rates in i*Xbp1* cells indeed indicated that ER-stressed IECs favored the full oxidation of 1C units to CO_2_ to generate NADPH (Fig. 2B+C). As expected, i*Xbp1* cells showed increased *Aldh1l2* expression at gene and protein level compared to iCtrl cells. Stimulation with TM also triggers *Aldh1l2* upregulation (Fig. 2D+S2B). ER-stress-dependent induction of *Aldh1l2* was also confirmed in intestinal organoid (Fig. 2E). Next, we analysed the gene expression status of additional 1C genes and observed that ER-stressed IECs predominantly increase the expression of the mitochondrial 1C genes *Shmt2*, *Mthfd2* and *Aldh1l2* whereas the expression of its cytosolic counterparts including *Aldh1l1* was unaffected (Fig. 2F).

**Fig. 2.**
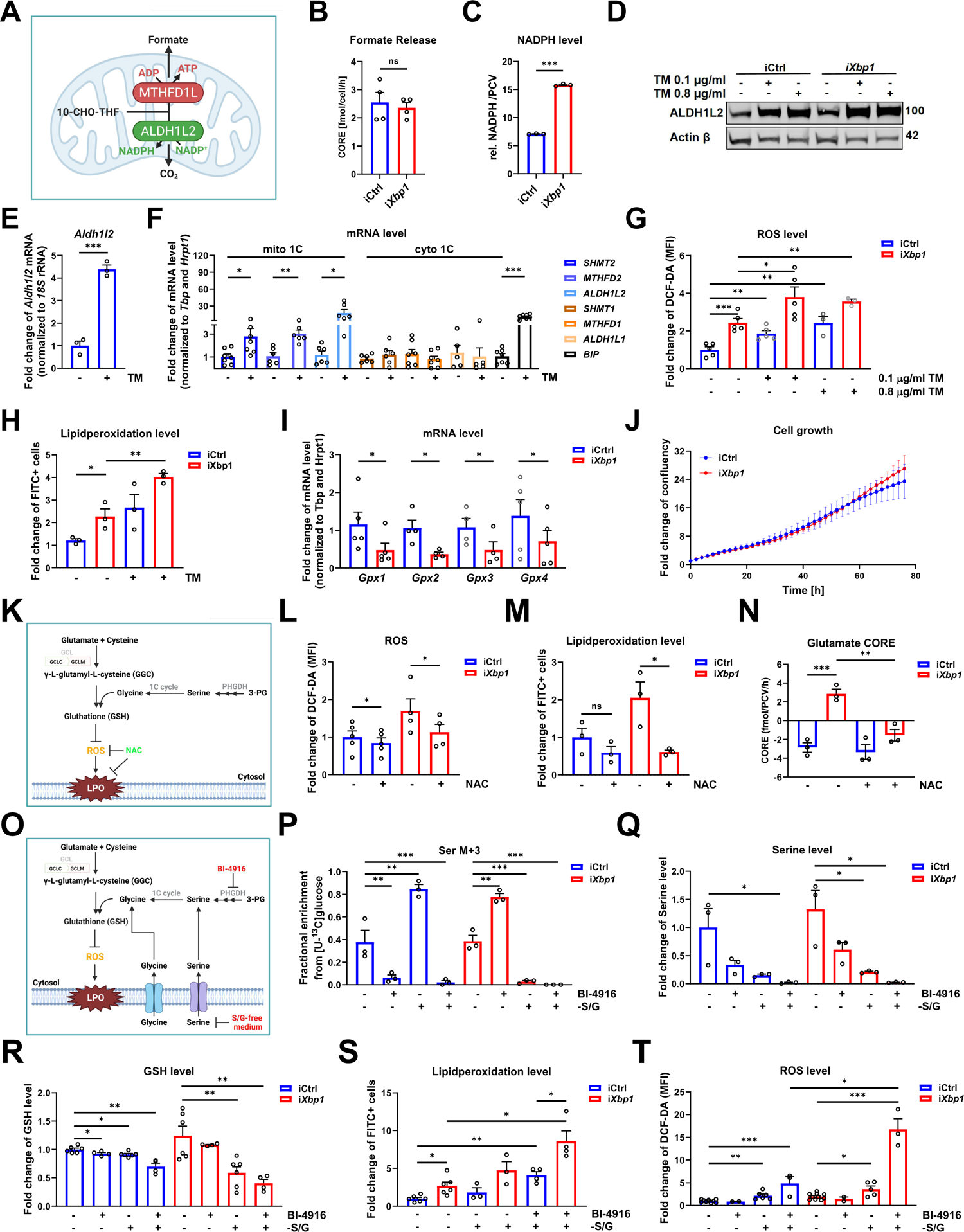
IECs activates the mitochondrial arm of the 1C cycle to counterbalance ER-stress-induced RedOx imbalance by boosting GSH and NADPH synthesis. (**A**) Schematic illustration of the 10-Formyl-Tetrahydrofolate (10-F-THF) branching point in the mitochondrial 1C metabolism (**B**) Absolute formate release rates (fmol/PCV/h) in iCtrl and i*Xbp1* cells (n = 4). (**C**) Relative NADPH level of iCtrl and i*Xbp1* cells (n = 3). (**D**) ALDH1L2 expression level in iCtrl and i*Xbp1* cells in the presence (+) and absence (-) of TM. VINCULIN serves as loading control. (**E**) *Aldh1l2* gene expression in murine SI organoids (derived from *Xbp1*^fl/fl^ mice) treated with 0.8 µg/ml TM for 24h (n = 3). (**F**) Relative mRNA expression level of mitochondrial (mito 1C) and cytoplasmic (cyto 1C) enzymes of the 1C cycle in wild-type Mode-K cells (iCtrl) treated with or without 0.8 µg/ml TM for 24h. ER chaperone *Bip*/*Grp78* serves as positive control for ER-stress induction. (**G**) Relative cytoplasmic ROS levels of iCtrl and i*Xbp1* cells treated with or without TM for 24h (n ≥ 3). (**H**) Relative percentage of FITC^+^ cells in iCtrl and i*Xbp1* cells treated 0.1 µg/ml TM for 24h (n=3). (**I**) Relative mRNA expression level of the indicated Gpx genes in iCtrl and i*Xbp1* cells (n=4, paired t-test). (**J**) Growth curve of iCtrl and i*Xbp1* cells (n = 3). (**K**). Schematic diagram of the antioxidant treatment strategy to reduce intracellular ROS level. (**L**) Relative cytoplasmic ROS level of NAC-treated iCtrl and i*Xbp1* cells (n ≥4, paired t-test). (**M**) Relative percentage of FITC^+^ cells in iCtrl and i*Xbp1* cells after 24h NAC treatment (n=3, paired t-test). (**N**) Absolute consumption and release rates of glutamate in iCtrl and i*Xbp1* cells after 24h treatment with 10 mM NAC (n ≥ 3). (**O**) Schematic illustration of the different intervention strategies used to interfere with GSH synthesis. (LPO = Lipid peroxidation) (**P**) Fractional enrichment of serine M+3 in iCtrl and i*Xbp1* cells cultured in complete or serine/glycine-free (-S/G) medium supplemented with 17 mM [U-^13^C] glucose tracer in the presence (+) and absence (-) of 15 mM BI-4916 for 24h (n=3, Two-Way ANOVA). (**Q**) Intracellular serine level of iCtrl and i*Xbp1* cells after 24h S/G starvation and co-treatment with 15 mM BI-4916 (n=3). (**R**) GSH level of iCtrl and i*Xbp1*cells; treated as in C (n ≥4). (**S**) Relative percentage of FITC^+^ cells in iCtrl and i*Xbp1* cells in complete or S/G-free medium and co-treatment with 15 mM BI-4916 (n≥ 3). (**T**) Relative cytoplasmic ROS level of iCtrl and i*Xbp1* cells after; treated as in C (n ≥2, paired t-test). \**P*<0.05, \*\**P* <0.01, \*\*\**P* <0.001.

Since the observed metabolic adaptations are crucial for maintaining cellular Redox homeostasis, we next checked the cytosolic ROS level. We noted increased ROS levels at baseline in i*Xbp1* cells which were further augmented in response to TM treatment (Fig. 2G). In line with increased ROS levels, LPO was also significantly elevated in i*Xbp1* cells and further increased upon TM stimulation (Fig. 2H). It was puzzling why the increase of GSH synthesis under chronic ER-stress does not lead to sufficient intracellular neutralisation of ROS and LPO species. Neutralisation of ROS is conducted via the reduction of hydroperoxides by glutathione peroxidases (GPXs) (*41*). We therefore tested the expression of *Gpx1-4* and observed significantly reduced mRNA levels of all *Gpx* genes in *Xbp1*-deficient cells (Fig. 2I). Probably, the reduced *Gpx* expression partially accounts for the increased accumulation of ROS in chronically ER-stressed cells. However, despite of increased Redox stress in *Xbp1* deficient cells, we did not observe any differences in proliferation, indicating that the metabolic adaptations prevent major oxidative damage (Fig. 1J). To test if metabolic adaptations in *iXbp1* cells are indeed the consequence of elevated ROS, we used ROS scavengers to rescue the metabolic phenotype (Fig.2K). Treatment with the antioxidant N-acetyl-cysteine (NAC) successfully lowered intracellular ROS and LPO levels (Fig. 2L+M). Most strikingly, NAC treatment also reverted the observed glutamate switch in i*Xbp1* cells (Fig. 2N). In sum, our findings illustrate that metabolic rewiring is directly linked to ER-stress-mediated ROS increase.

To test the importance of the observed metabolic polarisation in ER-stressed IECs, we targeted GSH homeostasis by limiting intracellular serine and glycine availability (Fig. 2O). We therefore deprived iCtrl/i*Xbp1* cells of serine/glycine (S/G) to limit GSH precursor availability. A decrease of intracellular serine and glycine levels as well as an accumulation of the precursor γ-L-glutamyl-L-cysteine (GGC) confirmed the effectiveness of the S/G starvation (Fig. S2C-E). However, under S/G deprivation, cells are able to activate the serine *de novo* synthesis pathway (SSP) to generate serine from glucose (*42*). To test for this compensatory mechanism, we performed [U-^13^C]-glucose tracing to monitor the relative glucose flux through the SSP. Illustrated by the M+3 serine isotopologue, S/G starvation strongly increased the flux through SSP. Of note, at standard conditions, SSP activity of i*Xbp1* cells was not increased compared to iCtrl cells (Fig. 2P). As the compensatory activation of SSP might directly feed into GSH synthesis, we combined S/G starvation with PHGDH inhibition, the rate limiting enzyme of the SSP pathway, by using the competitive inhibitor BI-4916 (*43, 44*). As expected, BI-4916 treatment blocked SSP activity and dual perturbation with BI-4916 and S/G starvation further reduced intracellular serine and GSH levels, confirming the efficacy of our treatment (Fig. 2P-R). Interestingly, GSH depletion was more pronounced in i*Xbp1* cells compared to iCtrl cells (Fig. 2R).

To assess whether the metabolic shift towards reduced GSH levels functionally mimicked the RedOx imbalance phenotype observed upon chronic ER-stress, we measured ROS and LPO levels in the presence of different interventions. Indeed, S/G deprivation with or without SSP inhibition led to increased LPO and ROS accumulation, which was again more pronounced in i*Xbp1* cells. In contrast to S/G deprivation, sole inhibition of SSP neither increased ROS nor diminished GSH level (Fig. 2R+T), indicating that the extracellular supply with S/G is the major source for GSH precursors and compensates for SSP loss in IECs. Although the cause for the impaired Redox homeostasis is slightly different, both chronic ER-stress as well as GSH limitation led to an inefficient removal of ROS and LPO species in IECs. Thus, our data point out that S/G supply is important to ensure proper RedOx homeostasis in ER-stressed IECs.

Overall, our data indicate that ER-stress induces a metabolic shift towards GSH metabolism to counterbalance ROS. Furthermore, we identify a distinct reprogramming of the 1C metabolism in response to ER-stress to ensure sufficient NADPH supply. Both metabolic adaptations are essentially needed to ensure a healthy Redox balance to sustain proliferation of IECs.

### Metabolic rewiring towards GSH and NADPH synthesis is of physiological relevance in IBD patients

To validate whether the metabolic adaptation of serine, glycine and 1C (SGOC) metabolism contributes to the pathogenesis of IBD, we analysed transcriptome dataset from UC and CD patients (*45*). Despite the low patient quantity, we observed a significant increase of *SLC7A11* and *ALDH1L2* expression in inflamed sigmoid colon biopsies of UC patients, and a similar trend in CD patients. In contrast, no expression changes were observed in non-inflamed UC patient as well as in samples from the healthy and disease control group (Fig. 3A).

**Fig. 3.**
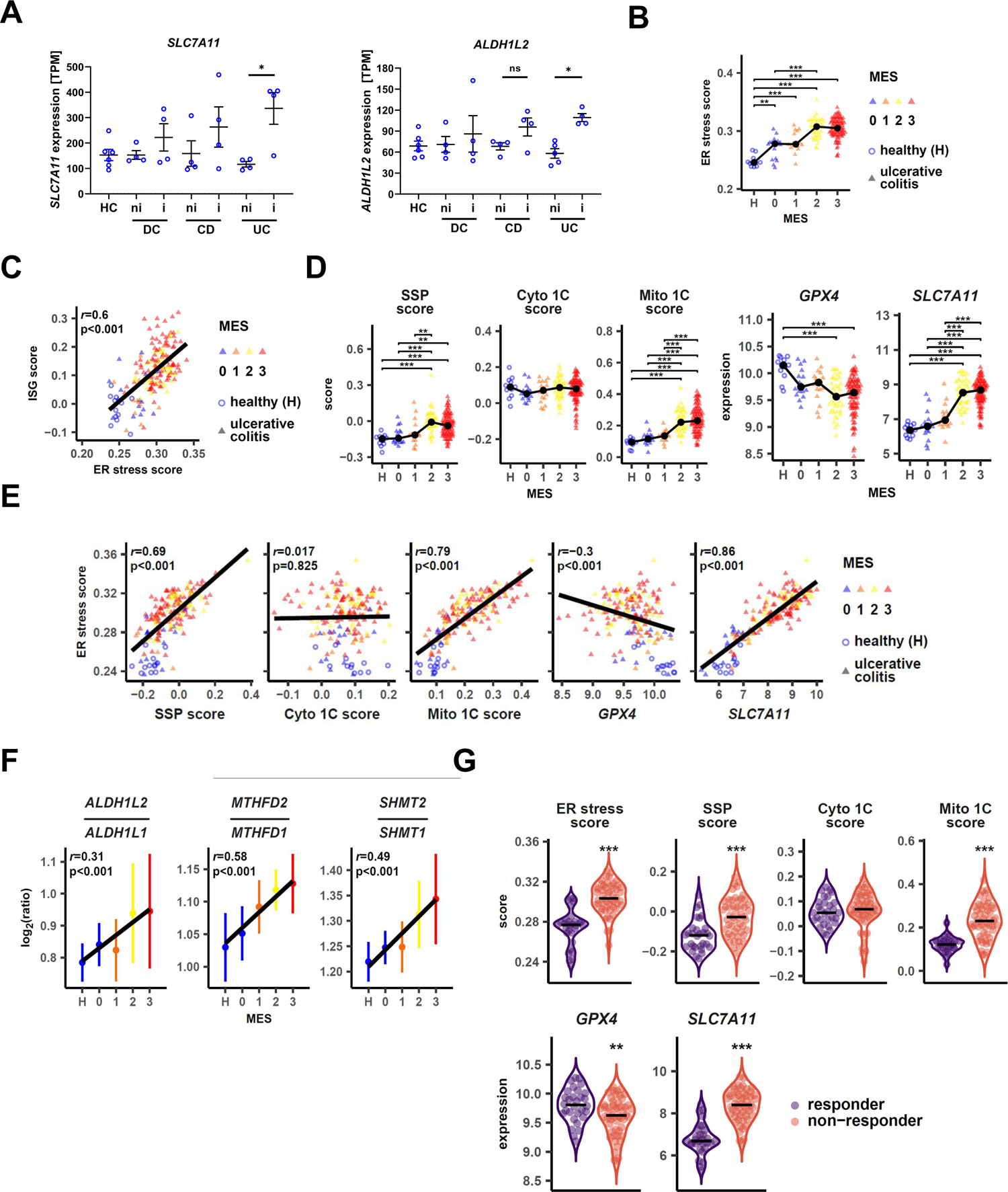
Metabolic adaption of the serine metabolism occurs in the inflamed intestinal epithelium of IBD patients. **(A)** RNAseq data analysis of *SLC7A11 and ALDH1L2* in patient samples derived from the sigmoid colon region. Samples derived from healthy donor (HC) or non-inflamed (ni) and inflamed (i) areas of Crohn’s (CD) or ulcerative colitis patients (UC). Inflamed and non-inflamed samples from an inflammatory disease unrelated to IBD serve as disease control (DC) (n ≥ 4 patients). (**B**) Correlation of ER-stress with endoscopic MAYO score (MES) in UC patients using the GSE73661 dataset and the singscore calculation method. ER-stress signature is defined by Powell *et al.* and includes 62 genes enriched in the colonic epithelial cells upon ER-stress (*12*). Black dots and connecting lines show the median expression. (**C**) Correlation of ER-stress with the interferon-stimulating gene signature (*31*) in the GSE73661 dataset. Pearson’s correlation coefficients *r* and p-values are shown. **(D**) SGOC metabolism related gene expressions and scores according to MES in UC patients from the GSE73661 dataset. Black dots and connecting lines show the median expression. (**E**) Correlation of SGOC metabolism-related gene expression with ER-stress and disease severity in the GSE73661 dataset. Pearson’s correlation coefficients *r* and p-values are shown. (**F**) Gene expression ratios of the mitochondrial 1C cycle enzymes over their cytosolic counterparts (*ALDH1L2*/*ALDH1L1*, *SHMT2/SHMT1*, *MTHFD2*/*MTHFD1* and their correlation with IBD severity. (**G**) Correlation of ER-stress and the expression of SGOC metabolism-related genes in patients from the dataset GSE73661 according to therapy response status \**P*<0.05, \*\**P* <0.01, \*\*\**P* <0.001.

To validate these results, we analysed two different transcriptome datasets from large UC patient cohorts (*46, 47*). We first tested the hypothesis whether ER-stress might contribute to the metabolic adaptations in the inflamed mucosa of UC patients. We therefore analysed the relation of mucosal ER-stress with the degree of inflammation by correlating disease severity with a previously published colon-specific ER-stress signature (*12*). We discovered a significant correlation of ER-stress with endoscopic and histologic disease severity (Fig. 3B+S3A). Additionally, we observed a strong correlation of ER-stress with an interferon-stimulated genes (ISG) signature (*31*) pointing towards a direct link between ER-stress and mucosal inflammation (Fig. 3C+S3B). We further delineated that the SSP including *PHGDH*, *PSAT1* and *PSPH* (SSP score) strongly correlated with disease severity (Fig. 3D+S3C) as well as the amplitude of ER-stress (Fig. 3E+S3D). Moreover, we also validated our *in vitro* findings and observed an increase in *SLC7A11* as well as a decrease in *GPX4* expression which was coupled to both parameters (Fig. 3D, 3E, S3C+S3D). Remarkably, the mitochondrial 1C genes score (Mito 1C, scoring *SHMT2*, *MTHFD2*, *ALDH1L2*) was also strongly linked to both parameters. In contrast, the cytosolic 1C genes score (Cyto 1C, scoring *SHMT1*, *MTHFD1* and *ALDH1L1*) did not show any dependency. By considering the gene expression ratios of the mitochondrial 1C enzymes over their cytosolic counterparts, we further support our *in vitro* findings in ER-stressed IECs. We observed a disease severity-dependent activation of the mitochondrial 1C metabolism in both transcriptome datasets (Fig. 3F and S3E). Additionally, colon biopsies of UC patients also showed an increased expression of the mitochondrial 1C enzymes compared to healthy controls (Fig. S3F) (*48*). Next, we checked the expression status of genes related to the SGOC metabolism in *Xbp1*-deficient mice as a model to mimic chronic ER stress in the gut (*5, 49*). Due to the limited sample number, we did not detect significant differences. However, we observed similar changes in regard to upregulation of mitochondrial 1C and SSP (Fig. S3G).

To further substantiate the hypothesis that the metabolic rewiring corresponds with disease activity in IBD, we assessed the differential changes of Mito1C and SPP signatures in UC patients under biologic therapy. Interestingly, the targeted treatment with biologic therapies led to a significant reduction of the metabolic signature in therapy-responding patients, suggesting that the metabolic remodelling is an important protective cellular adaptation to counterbalance chronic inflammation (Fig. 3G).

As our data collectively demonstrate that a metabolic rewiring of the SGOC metabolism takes place in IBD patients, we furthermore aimed to understand whether this is a cell type-specific adaptation to ER-stress. We therefore evaluated single-cell RNA sequencing (scRNA-Seq) data obtained from the SCP259 scRNA-Seq IBD dataset (colon mucosa of 18 UC patients and 12 healthy individuals (*50*)). We observed in different epithelial cell clusters an upregulation of SSP and mitochondrial 1C genes in a disease activity-dependent manner which correlates with the amplitude of the ER-stress signature (Fig. S3H+I). Interestingly, this metabolic adaptation was predominantly found in clusters of secretory epithelial cells (secretory TA, immature goblet cells) and transient amplifying (TA) cells, stem cells, which are, due to their proliferative and secretory demand highly dependent on effective endoplasmic reticulum function. In summary, transcriptome data from UC patients further corroborate a crucial role of metabolic reprogramming of 1C metabolism in the context of intestinal inflammation.

### Chronic ER-stress-induced RedOx imbalance leads to cGAS/STING-signalling exhaustion

Having established the importance of 1C metabolism in the intestinal epithelium, we wondered whether ER-stress induced alterations in the 1C metabolism affect innate immune pathways relevant for IBD. It is widely accepted that disease extent and disease severity of IBD patients directly correlate with an increased risk of viral infection such as CMV (*28, 51*), representing a major clinical challenge in the disease management. As CMV control critically depends on cGAS/STING dependent IFN-production (*52*), and both ROS and by-products of the LPO reactions (e.g. 4-HNE) suppress cGAS/STING-signalling by either preventing STING dimerisation or its transition from the ER to the Golgi compartment (Fig. 4A) (*21, 53*), we tested a potential crosstalk between the observed ER-stress-induced metabolic imbalance and cGAS/STING-signalling in IECs.

**Fig. 4:**
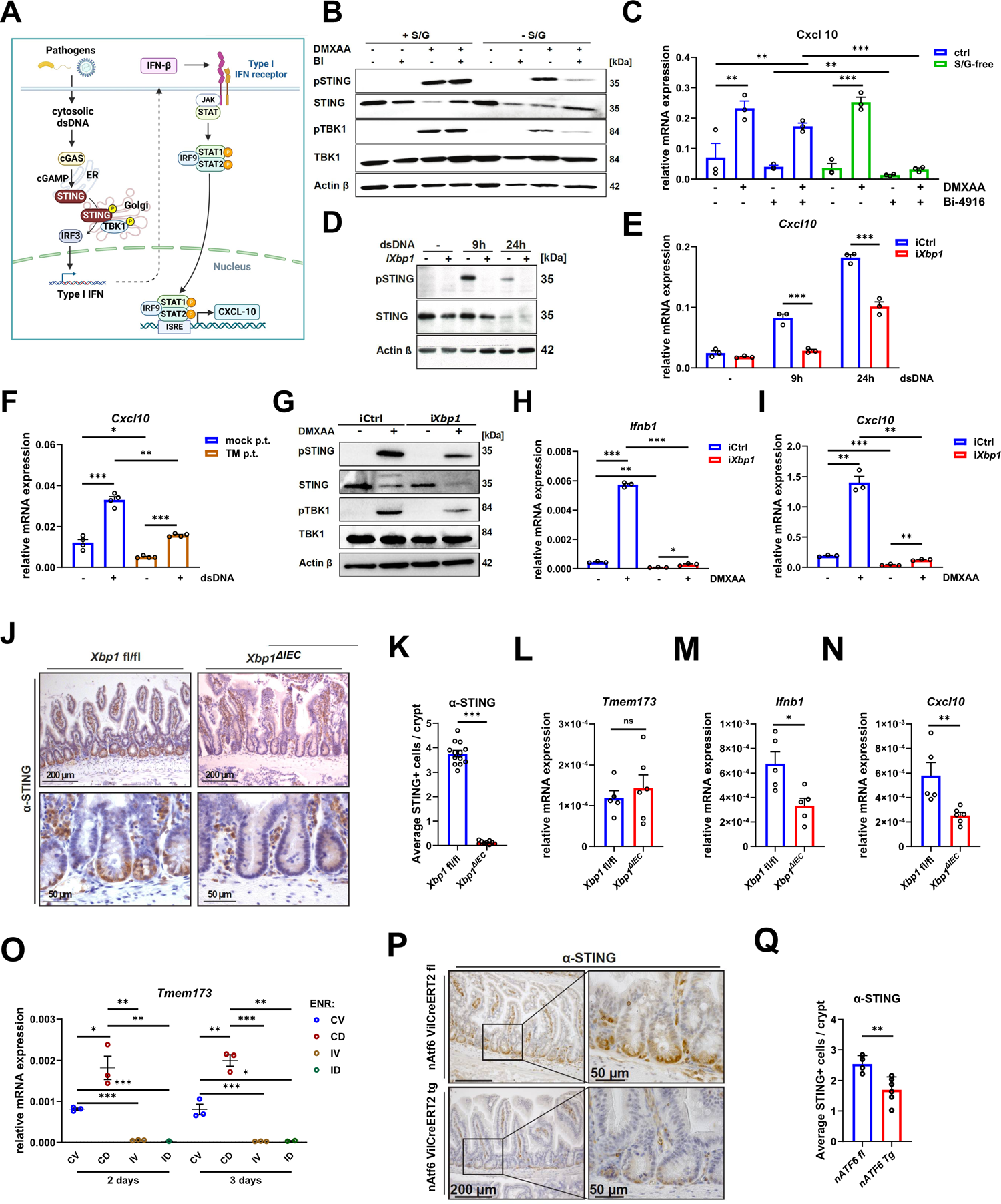
Chronic ER-stress-induced RedOx imbalance dampens cGAS/STING-signalling *in vitro* and *in vivo*. (**A**) Schematic outline of the cGAS/STING-signalling and its downstream targets. (**B**) Western Blot analysis of iCtrl cells stimulated with DMXAA (100 µg/ml, 1h) after cultivation in complete or S/G-free medium for 24h and co-treatment with 15 mM BI-4916 (**C**) *Cxcl10* gene expression of iCtrl and i*Xbp1* cells which were starved for S/G and treated with 15 μM BI-4916 for 24h before stimulation with DMXAA (100 μg/ml, 3h) (n=3). (**D**) Western Blot analysis of STING activation in iCtrl and i*Xbp1* cells stimulated with dsDNA (1 µg/ml) for the indicated time points. (**E**) *Cxcl10* gene expression of the experiment shown in (D) (n=3). (**F**) *Cxcl10* gene expression in dsRNA-stimulated (1 µg/ml) iCtrl cells pretreated with or without 0.1 µg/ml Tunicamycin (n=4). (**G**) iCtrl and i*Xbp1* cells were stimulated with DMXAA (100 µg/ml, 1h) and expression of the indicated proteins assessed by Western Blot. (**H+I**) *Ifnb1* (H) and *Cxcl10* (I) gene expression of iCtrl and i*Xbp1* cells stimulated with DMXAA (100 µg/ml, 3h, n=3). (**J**) IHC staining of STING expression in *Xbp1*^fl/fl^ (n=13) and *Xbp1*^ΔIEC^ (n=10) mouse SI sections. Representative images. (**K**) Corresponding quantification of STING-positive cells per crypt. (**L-N**) *Tmem173* (L), *Ifnb1* (M) and *Cxcl10* (N) gene expression from SI crypts isolated from *Xbp1*^fl/fl^ (n= 5) and *Xbp1*^ΔIEC^ (n= 6) mice. (**O**) *Tmem173* gene expression in murine SI organoid cultures enriched for stem cells (CV), Paneth cells (CD), enterocytes (IV) and Goblet cells (ID) after cultivation in specific enrichment media (ENR) for 2d or 3d. (**P**) IHC staining of STING expression in *nATF6* VilCreERT2 tg or fl mice. Intestinal epithelial-specific overexpression of ATF6 was induced with Tamoxifen for 7d and sacrificed after another 4d. Representative HE/STING-stained SI sections. (**Q**) Corresponding quantification of STING-positive cells per crypt (n=4 (fl) and n=6 (tg)).*p<0.05, **p<0.01, ***p<0.001.

Since our previous experiments indicate that S/G deficiency as well as chronic ER-stress causes an accumulation of ROS and LPO species, we asked whether S/G deprivation disrupts epithelial STING signalling. We limited S/G availability by either the single (S/G starvation) or double intervention (S/G starvation + PHGDHi) and analysed STING activation in iCtrl cells in the presence or absence of the STING agonist DMXAA. While S/G starvation or BI-4916 treatment alone had no significant impact on STING activation, dual perturbation resulted in decreased phosphorylation of STING and its downstream target TBK1 (Fig. 4B). Additionally, a strong and significant reduction of *Cxcl10* expression was observed following the dual treatment (Fig. 4C). Altogether, our data show that the supply of serine and glycine is critical to facilitate proper STING/IFN-I signalling in ER-stressed IECs.

As *Xbp1* deficiency also induces RedOx imbalance in a GSH-dependent manner, we postulated that chronic ER-stress impairs IFN response to viral infection. To test this hypothesis, we stimulated iCtrl and i*Xbp1* cells with double-stranded (ds) DNA and noticed a strong abrogation of STING activity (Fig. 4D) and *Cxcl10* expression (Fig. 4E) in *Xbp1*-deficient IECs. This observation was phenocopied by treating wild-type cells with the chemical ER-stress inducer TM prior to dsDNA exposure (Fig. 4F). Our findings suggest that unresolved ER-stress promoted exhaustion of the STING pathway with impaired IFN-I and CXCL10 production in response to pathogen sensing. Indeed, the STING agonist DMXAA activated STING and TBK1 phosphorylation in iCtrl cells whereas the effect was reduced in i*Xbp1* cells (Fig. 4G). This finding was accompanied by impaired *Ifnb1* and *Cxcl10* induction (Fig. 4H+4I). Collectively, our data support the hypothesis that metabolic reprogramming of the 1C metabolism upon ER-stress directs STING signalling in the intestinal epithelium.

Next, we aimed to understand the *in vivo* consequences of chronic ER-stress on cGAS/STING-signalling. We thereby took advantage of an intestinal-specific conditional *Xbp1* knockout mouse model and analysed STING expression as well as IFN-I-activity (*5, 54*). We noted that *Xbp1*^ΔIEC^ mice showed significantly reduced STING expression compared to *Xbp1*^fl/fl^ (wiltype) littermates (Fig. 4J). Especially the crypt-specific STING expression was strongly abolished upon deletion of *Xbp1* (Fig. 4K+S4A). Notably, reduced STING protein expression was unrelated to *Sting* transcription (Fig. 4L), indicating post-translational regulation. Additionally, we observed reduced *Cxcl10* and *Ifnb1* expression in small intestinal (SI) crypts from *Xbp1*^ΔIEC^ mice, an indication for abrogated STING/IFN-I activity (Fig. 4M+N). Lastly, we co-validated the impact of prolonged ER-stress on epithelial cGAS/STING signalling using an independent transgenic mouse model, i.e. *nAtf6*^IEC^ VilCreERT2 mice carrying a Tamoxifen-inducible intestinal epithelial *Atf6* overexpression (leading to constitutive ER-stress (*55, 56*)). Tamoxifen-induced overexpression of *Atf6* also evoked epithelial STING suppression (Fig. 4P+Q).

To reveal which specific cell-type predominantly expresses STING in the intestinal epithelium, we subjected SI organoids to different conditioned media to enrich for specific epithelial lineages, i.e. stem cells, Paneth cells, Goblet cells and enterocytes (Fig. 4O) (*57*). Lysozyme expression confirmed the SI organoid differentiation into the different epithelial lineages (Fig. S4B+C). Notably, *Tmem173* expression was most pronounced in Paneth cell-enriched organoids (ENR-CD) and stem cell-enriched organoids (ENR-CV), while enrichment for enterocytes (ENR-IV) or goblet cells (ENR-ID) did not show *Tmem173* expression (Fig. 4O). The finding that Paneth cells predominantly express STING and that *Xbp1* loss is known to trigger increased Paneth cell death in intestinal crypts (*5*) further corroborates a crucial connection between ER-stress and mitochondrial 1C metabolism on cGAS/STING signalling.

### Chronic ER-stress increases susceptibility to bacterial and viral infections

To test whether ER-stress-induced perturbation of STING signalling links to susceptibility to viral or bacterial infections, we infected i*Xbp1* cells (and controls) with murine CMV (MCMV) and assessed STING-dependent IFN-I production. Indeed, iCtrl cells responded to MCMV infection with STING activation, IFN-I induction and subsequent viral containment (Fig. 5A-C). In contrast, *Xbp1*-deficient cells showed an increased MCMV load and failed to induce *Ifn1b* and *Cxcl10* expression. To verify the impact of epithelial STING signalling on CMV infection *in-vivo*, we challenged *Xbp1*^fl/fl^ and *Xbp1*^ΔIEC^ mice with MCMV intraperitoneally. Five days after initial infection, we measured the degree of intestinal inflammation, as described previously (*58, 59*). In line with previous reports, total viral load within the intestinal mucosa was not detectable at day of sacrifice. However, we observed increased severity of enteritis by a semi-quantitative histology score in *Xbp1*^ΔIEC^, but not in *Xbp1*^fl/fl^ mice (Fig. 5D+E). This observation confirmed that MCMV infection controlled the inflammatory tone in genetically susceptible mice that exhibit chronic epithelial ER-stress. Finally, we infected IECs with the intracellular pathogen *Listeria monocytogenes* to test whether the observed STING exhaustion is also relevant to bacterial infection. As described before (*60*), *L. monocytogenes* infection resulted in strong *Cxcl10* and *Ifnb1* induction in iCtrl cells, which was barely detectable in i*Xbp1* cells (Fig. 5F+5G). Collectively, our data indicate that ER-stress causes a functional impairment of STING signalling which leads to inefficient pathogen clearance in the intestinal epithelium.

**Fig 5:**
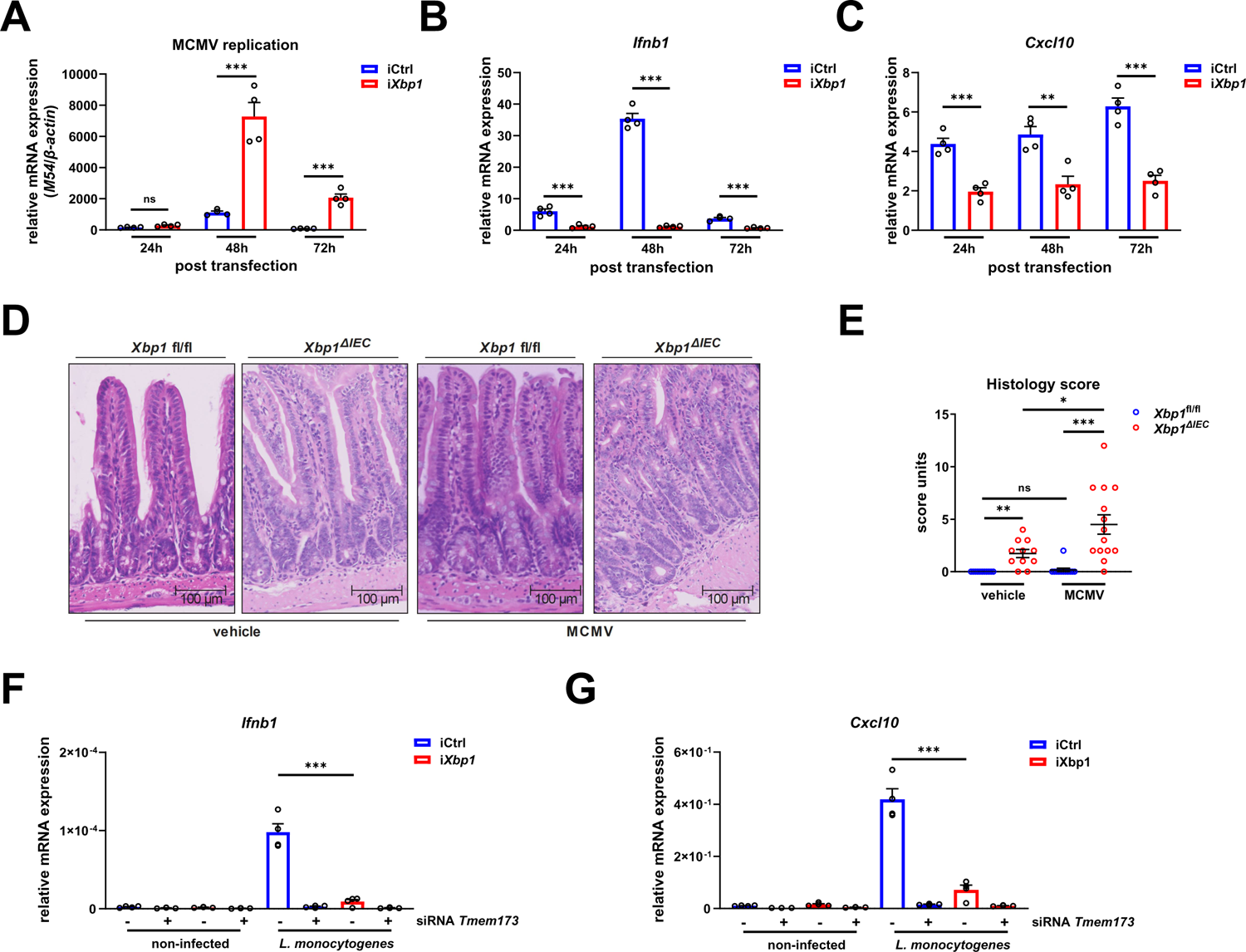
Chronic ER-stress increases susceptibility to bacterial and viral infections. (**A**-**C**) iCtrl and i*Xbp1* cells were infected with MCMV at a MOI of 10 and MCMC viral load (A), *Ifnb1* (B) and *Cxcl10* (C) gene expression was assessed by qRT-PCR (n= 4). (**D+E**) *Xbp1*^fl/fl^ and *Xbp1*^ΔIEC^ mice were infected intraperitoneally with vehicle (n=11 for each group) or MCMV (n=12 (*Xbp1* fl/fl); n=14 (*Xbp1*^ΔIEC^)) and intestinal inflammation was assessed via histology. Representative images of HE stained SI sections (D) and quantification of the histology score (E) are illustrated. (**F+G**) iCtrl or i*Xbp1* cells in the presence (+) or absence (-) of *Tmem173* were exposed to *Listeria monocytogenes* at a MOI of 50 for 1h before medium was exchanged and Gentamicin (50 µg/ml) was added for 1h, followed by Gentamicin at 5 µg/ml for 22 h. STING activation was monitored via the *Ifnb1* (**F**) and *Cxcl10* (**G**) gene expression (n=4). *p<0.05, **p<0.01, ***p<0.001.

### Antioxidant therapy restores cGAS/STING-signalling and CMV control in IECs

After demonstrating that antioxidant treatment alleviates the ER-stress-mediated metabolic adaptation in IECs, we finally tested the hypothesis that NAC supplementation restores epithelial STING signalling and thus protects against CMV infections. We observed that antioxidant treatment restored DMXAA-induced STING and TBK1 phosphorylation in i*Xbp1* cells (Fig. 6A), showing that ER-stress indeed impairs STING signalling via ROS and/or LPO generation. Alternatively, GPX4 inhibition, using the GPX4 inhibitor RSL3, led to a nearly complete abrogation of STING signalling in iCrtl cells, which was rescued by NAC treatment (Fig. S5A+B). Similarly, the impact of combined S/G starvation and PHGDH inhibition on STING signalling was partially neutralised in the presence of NAC (Fig. 6B+C). Finally, we wondered whether restoration of STING signalling via NAC treatment directly improves CMV infection control in IECs. To this end, we co-treated MCMV infected iCtrl and i*Xbp1* cells with the antioxidant. In line with our hypothesis, i*Xbp1* cells showed a robust restoration of IFN-I signalling (Fig. 6D+E) which further resulted in reconstituted suppression of MCMV viral load (Fig. 6F). Collectively, our data suggest that ER-stress-derived ROS increase limits STING signalling, representing the underlying reason for impaired viral control.

**Fig 6:**
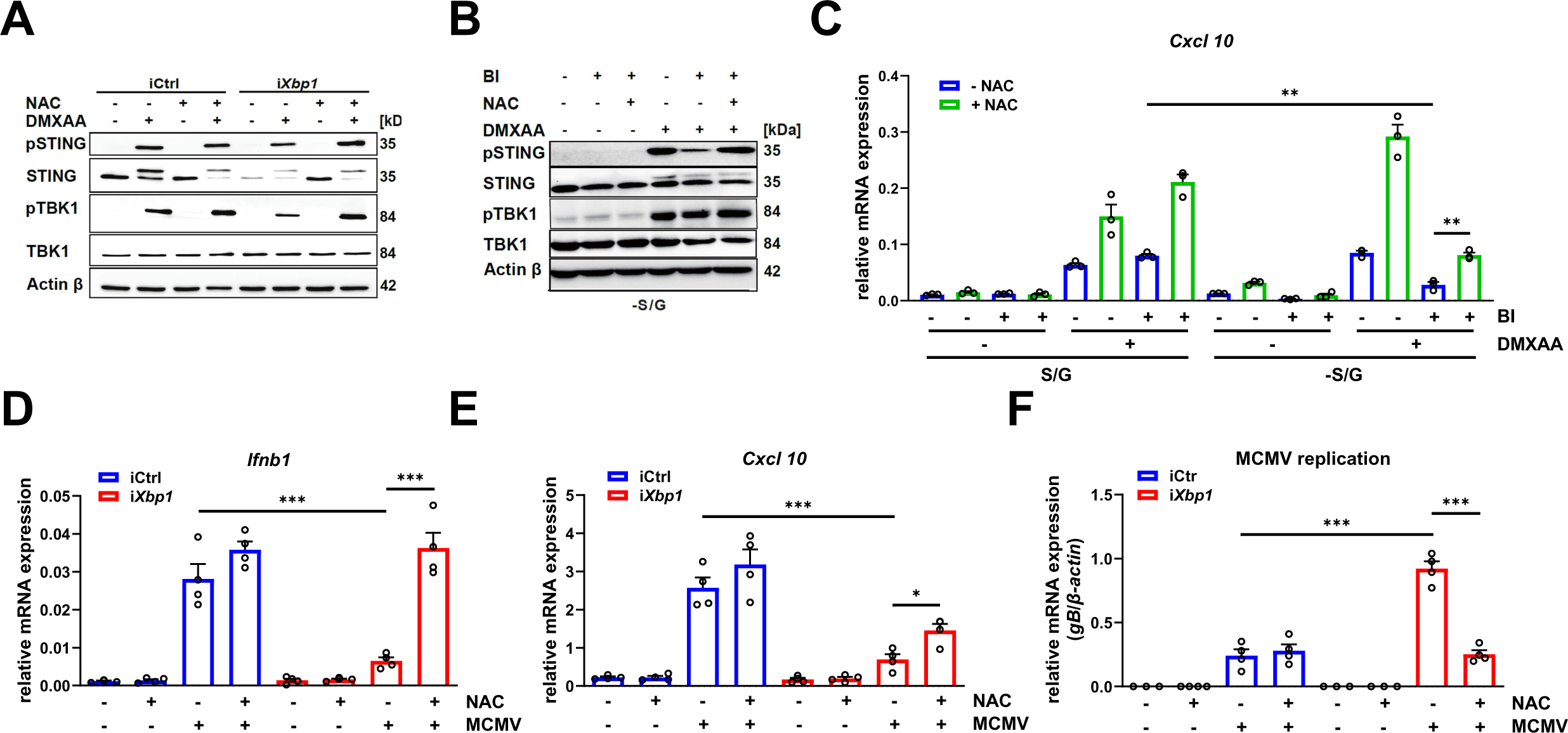
Antioxidant therapy restores cGAS/STING signalling and improves CMV infection control in intestinal epithelial cells. (**A**) Western Blot analysis of iCtrl and i*Xbp1* cells stimulated with DMXAA (100 μg/ml, 1h) after 24h pretreatment with 10 mM NAC. (**B**) Western blot analysis of iCtrl cells stimulated with DMXAA (100 µg/ml, 1h) after cultivation in complete or S/G-free medium for 24h in the presence or absence of 10 mM NAC and 15 mM BI-4916. (**C**) *Cxcl10* gene expression of iCtrl stimulated with DMXAA (100 μg/ml, 3h) after 24h S/G starvation and co-treatment with 10 mM NAC and 15 mM BI-4916 (n=3).(**D-F**) iCtrl and i*Xbp1* cells were stimulated with 10 mM NAC and infected with MCMV (MOI = 1). mRNA expression of *Ifnb1* (H), *Cxcl10* (I) as well as viral load (J) (measured by viral DNA copy number) were determined via qRT-PCR (n=4). *p<0.05, **p<0.01, ***p<0.001.

## Discussion

The intestinal epithelium is not only critical for nutrient and water absorption from food but also forms a tremendous protective barrier against the outer environment (*61*). Due to the frequent exposure to extrinsic and intrinsic stress stimuli (e.g. nutrient availability, harmful pathogens, microbiota-derived metabolites etc.), cells have to constantly adapt their cellular and metabolic programs to maintain intracellular homeostasis (*62*). Although prior studies already showed a direct link between the manifestation of chronic ER-stress and the development of IBD (*32, 63*), there is a limited understanding of how IECs fine-tune their metabolism to deal with unresolved ER-stress and related gut inflammation.

Here we demonstrate that chronically ER-stressed IECs activate a distinct metabolic program to boost GSH synthesis. We thereby observed that IECs actively increase the intracellular availability of amino acids especially required for GSH synthesis by modulating amino acid transporter activity in the plasma membrane. For example, upregulating of the SLC7A11 transporter system or increased glutamine uptake in response to chronic ER-stress are two options how IECs cover their additional cysteine and glutamate demands for GSH synthesis. Interestingly, the observed glycine switch is supported by data from human IECs. Howard *et al*. demonstrated that the glycine transporter SLC6A9 (Glyt1) is crucial to maintain intracellular GSH level upon increased oxidative stress (*64*). Previous data indeed indicate that the fine-tuning of amino acid transporters including SLC6A9 and SLC7A11 is one common strategy to cover the specific amino acid demands upon ER-stress (*65*). Surprisingly, depending on the ER-stress amplitude, IECs seems to coordinate the support of GSH synthesis differently. Under chronic ER-stress, IECs utilise serine as main source to cover cellular glycine demands. In contrast, the direct uptake of extracellular glycine is favored upon acute ER-stress conditions. In this case, the additional glycine demand for GSH synthesis most likely exceeds the maximal capacity of the SSP to provide precursors for both GSH and purine synthesis (*66*). Interestingly, the dependency on extracellular glycine is presumably compensated by the long-term metabolic and cellular adaptations in chronically ER-stressed IECs.

We further showed that IECs induce a second compensatory mechanism by upregulating the mitochondrial arm of 1C cycle towards ALDH1L2. Thus, IECs are an excellent showcase for the proposed metabolic flexibility of the 1C metabolism and shows how ER-stressed cells maximise their supply for GSH synthesis under ER-stress conditions (*16*). Firstly, the SHMT2 reaction provides additional glycine as building block for GSH. Secondly, ALDH1L2 upregulation increases NADPH supply. Since the mitochondrial and cytosolic NADPH pools are separated from each other and NADPH shuttling does not occur (*67*), the additional NADPH presumably boosts the conversion of mitochondrial GSSG to GSH. Several studies already showed that ALDH1L2 is of great importance to regulate mitochondrial ROS level (*68–71*). However, we cannot exclude that other metabolic sinks for ALDH1L2-derived NADPH exist in ER-stressed IECs. For example, under normal growth conditions, ALDH1L2-derived NADPH is known to support proline biosynthesis (*71*).

Although the mechanistic basis for the metabolic reprogramming in IECs deserves to be resolved in more detail, recent studies suggest a critical role for the PERK-ATF4-pathway and the integrated stress response (ISR) (*72–74*). CHIPseq data already demonstrated that *PHGDH*, *PSAT1*, *SHMT2, MTHFD2* and *ALDH1L2* are direct ATF4 targets (*75*). Additionally, several amino acid transporters including *SLC7A11*, *SLC6A9* and the serine transporters *SLC1A4* as well as *SLC1A5* possess ATF4 binding sites and are upregulated in response to ER-stress in a PERK-dependent manner (*65, 75*). Along the same line, both deletion of ATF4 or ISR inhibition prevent the transcriptional upregulation of *ALDH1L2*, *SLC7A11* and *SLC6A9* in FH- and SDH-deficient cells (*74*). Interestingly, recent data demonstrate that glucose limitation also causes an upregulation of the mitochondrial 1C cycle towards *ALDH1L2* whereat the underlying mechanism(s) are not completely explored and might also be related to increased ER-stress (*38, 76*).

Paradoxically, although both metabolic adaptations are designed to increase the antioxidant capacity of IECs, chronic ER-stress causes a state of RedOx imbalance. Based on our findings, we postulate that IECs remodel their amino acid and 1C metabolism to accelerate the detoxification of ROS species upon ER-stress. Due to this metabolic adaptation, IECs normally receive a short-term buffer to activate compensatory mechanism(s) (e.g. UPR activation, increase of ER folding capacity etc.) and to restore cellular homeostasis. If ER-stress is not resolved in this initial phase or malfunctions of the compensatory systems (e.g. UPR) occurs, the rising ROS burden overwhelms the cellular detoxification systems and disrupts the RedOx balance. Despite of increased ROS, the metabolic reprogramming is still sufficient to guarantee the survival of Xbp1-deficient IECs indicated by their capability to maintain growth rates similar to wild-type cells. Since prolonged ER-stress is known to inhibit global protein translation (*77*), it is not surprising that several glutathione peroxidases are less expressed, giving a possible explanation why Xbp1-deficient IECs fail to completely restore RedOx homeostasis in the long run. Interestingly, the GSH-dependent glutathione peroxidase GPX4 is essential for the conversion of harmful lipid hydroperoxides into non-toxic lipid alcohols (*78*), partially explaining the increased LPO level observed in our chronic ER-stress model. Intriguingly, S/G starvation also leads to an increase in ROS and LPO. In contrast to the reduced GPXs activity upon chronic ER-stress, GSH limitation is the driving force for a deregulated RedOx balance upon S/G deprivation. Both chronic ER-stress as well as S/G starvation increase the GSH dependence of IECs and links our metabolic phenotype to GSH metabolism. Notably, our rescue experiments with the antioxidant NAC further underpin our notion that the metabolic rewiring represents a compensatory response to ER-stress and is coupled to an increase in ROS.

Importantly, our computational analysis demonstrates the *in vivo* relevance of the metabolic compensatory program in a clinically-relevant disease context. We directly linked the metabolic phenotype to the amplitude of ER-stress, the degree of inflammation as well as the disease severity status of UC patients. Beside the activation of the mitochondrial 1C cycle and SSP (also shown in Xbp1-deficient mice), UC patients also showed *SLC7A11* upregulation and *GPX4* downregulation in an ER-stress-dependent manner. Interestingly, increased LPO levels and reduced GPX4 activity are already reported in Crohn’s disease (*79*), supporting our notion that our findings are also relevant for CD patients. Our single cell analysis further reveal a cell-type specificity. Especially transient amplifying (TA) stem and immature goblet cells show the metabolic adaptation, whereas other cell types (e.g. enterocytes, M cells) presumably rely on SGOC-independent mechanisms to cope with increased ER-stress. Another interesting observation in TA cells is that increased mitochondrial activity is accompanied with reduced cytosolic 1C activity. Previous studies already reported that UPR defects induce a hyper-proliferative IEC phenotype in mice (*5*) and that UPR activation is described as an important mediator for the transition of stem cells into TA cells (*80*). Therefore, it would be interesting to check whether the metabolic rewiring of the SGOC is not only crucial for RedOx control but might also represent an important gatekeeper to maintain a healthy epithelial lining and to prevent uncontrolled crypt growth. From the clinical point of view, the fact that the metabolic signature is strongly linked to both disease severity and therapy-response could be exploited to define severity grade and/or therapy-response status of UC patients in the future.

Our finding that ER-stress-mediated impairment of the cGAS-STING signalling increases the susceptibility towards CMV infection is of great clinical importance. Mucosal CMV infections are a leading cause of complicated disease course in IBD (*81, 82*). Up to now, the underlying molecular pathophysiology driving increased susceptibility towards CMV colitis in IBD patients are poorly understood. In this study, our data provide a novel molecular nexus on how chronic ER-stress paves the susceptibility to CMV infection and thereby generate a molecular inside into a well-established and highly relevant disease complication in IBD. We thereby found first evidences that chronic ER-stress-caused RedOx imbalance prevents an efficient ROS/LPO clearance which inhibits cGAS-STING signalling. Our hypothesis is further underpined by our antioxidant rescue results and are in line with other studies demonstrating that both ROS and LPO by-products are able to disrupt STING function (*21, 53*). However, our *in vivo* results related to Paneth cells suggest that additional cell-type specific consequences of ER-stress might contribute to the overall impaired intestinal cGAS-STING-response. With regard to our antioxidant experiments, several studies demonstrated that ROS inhibition suppresses viral replication. Multiple herpesviruses are able to promote intracellular ROS generation and thereby support their own replication (*83*). In this context, it is shown that NAC supplementation inhibits the lytic replication cycle of Kaposi’s sarcoma-associated herpesvirus *in vivo* (*84*). Additionally, MHC68 replication is also suppressed in NAC-treated mice in a STING-dependent manner (*53*), pointing towards a direct link between ER-stress induced ROS increase, impaired cGAS-STING signalling and viral replication. We therefore postulate that the metabolic reprogramming has a profound impact on host immunity and might provide a layer of susceptibility towards pathogen infection in the inflamed mucosa of IBD patients. Recent data indeed indicate that the intestinal epithelium represents the primary sensor of mucosal CMV infection (46). Overall, our data strongly put forward the idea of an ER-stress driven immune exhaustion of the cGAS-STING pathway in the intestinal epithelium as a critical layer of susceptibility for CMV infection in IBD. Thus, our findings might pave the way towards future healthcare concepts to increase the viral control in IBD patients.

### Limitations of the study

While our S/G starvation experiments provide evidence that IECs remodel their GSH metabolism in response to ER-stress to sustain cGAS-STING-signalling, we did not exhaustively test the impact of the mitochondrial 1C cycle (especially ALDH1L2) on this viral defence pathway. Additionally, we showed that our findings regarding the metabolic phenotype were also relevant in the context of IBD. However, we exclusively focused on UC patients cohorts, even though CD patients show signs for a comparable metabolic remodelling towards GSH and 1C metabolism. Although we demonstrate that chronic ER-stress impairs STING signalling and CMV control in different murine models, our *in vivo* work is limited. In the future, it would be desirable to investigate the *in vivo* consequences of SG limitation on cGAS-STING-signalling in murine models in more detail. In this study, our infection models were solely based on murine CMV due to species-specific replication restrictions (*85*). Thus, the direct translation of all our findings to human CMV may be difficult without further investigation. Since direct *in vivo* experiments with human CMV are difficult, the usage of humanised mouse models could represent an alternative strategy to confirm our main findings in the future (*86*).

## Materials and Methods

### Experimental design

#### Mice

*Xbp1*^fl/fl^ mice were generated by targeted insertion of LoxP sites flanking the exon 2 of the *Xbp1* gene. VillinCre+/− mice (strain: B6.SJL-Tg (Vil-cre)997Gum/J) were purchased from Jackson Laboratory. Conditional knockout of *Xbp1* in the intestinal epithelium was then established by crossing VillinCre+/− mice with *Xbp1*^fl/fl^ mice, resulting in specific deletion of *Xbp1* (*Xbp1*^ΔIEC^) in the intestinal epithelium (*87*). Tamoxifen-inducible nATF6-HA overexpressing Villin-Cre n*Atf6*^IEC^ mice (n*Atf6* VilCreERT2 transgene (tg)) mice were generated as described previously (*55*). Overexpression of nATF6-HA was induced using Tamoxifen for seven days, and mice were sacrificed after another 4d allowing for full induction of overexpression. *Tmem173*^-/-^ mice were purchased from Jackson Laboratory (previously described by Jin et al., 2011, strain #025805) (*88*). Mice were housed under specific pathogen-free (SPF) conditions in individually ventilated cages (IVCs). Animals were held in a 12h light-dark cycle at 21°C ± 2°C and 60% ± 5% humidity. Food and water were available ad libitum. All animal experiments were approved by federal authorities and performed in accordance to institutional guidelines.

### Cell lines

In this study, the SV40 large T-antigen-immortalised murine small intestinal epithelial cell line Mode-K (51) was employed as an *in vitro* model for the intestinal epithelium. Mode-K cells harbouring an RNA interference (RNAi) based knockdown of *Xbp1* and a control RNA interference (*iXbp1*, iCtrl) were kindly gifted by Artur Kaser, University of Cambridge (6). In brief, RNA interference was achieved by stable introduction of a small hairpin RNA specific for *Xbp1* and a control small hairpin RNA identical to (*89*), except that SFGΔU3hygro was used. Knockdown of *Xbp1* was confirmed by qRT-PCR.

### Organoid generation and handling

Mouse organoids were cultivated as described previously (*57, 90*). To generate sustainable cultures of small intestinal organoids, mice were sacrificed and the small intestine was removed and cut into pieces. PBS wit 10 nM EDTA was added to the pieces for 10 min and the specimen was placed under mild agitation. Afterwards, the supernatant was removed, fresh PBS/EDTA solution was added and incubated under mild agitation for 10 min again. This was repeated four times. To remove debris, the suspension was filtered through a 100 µm strainer and centrifuged at 1200 rpm for 10 min. Crypts were then either used for protein respectively RNA isolation or resuspended in Matrigel and embedded in 24-well plates to generate organoid cultures. Medium was changed twice per week. Organoids were stimulated after 5 to 7 days of cultivation.

### Chemicals and siRNA-mediated knockdown

The following chemical stimulants were used (name, company, catalogue-number): DMXAA/Vadimezan (Hölzel, HY-10964), N Acetyl-L-cystein (Th. Geyer/Sigma, A7250-25G), BI-4916 (Hoelzel, HY-126253-5mg), RSL3 (Selleck Chemicals, S8155). siRNA knockdown of *Tmem173* was performed using the Viromer® Blue kit (Lipocalyx, Halle, Germany) according to the manufacturer’s protocol. *Tmem173* siRNA was purchased from Qiagen (catalogue-number GS72512).

### Cell culture

Mode-K cells were cultured in Dulbecco’s modified Eagle’s medium (DMEM) without phenol red, glucose, and glutamine (Thermo Fisher Scientific) and supplemented with 2 mM glutamine, 17 mM glucose, and 10% fetal bovine serum (FBS) at 37°C and 5% CO_2_. For serine/glycine starvation experiments cells were kept in customised DMEM medium (Thermo Fisher Scientific) supplemented with or without 400 μM glycine and serine, 2 mM glutamine, 17 mM glucose and 10% dialysed FBS.

### *In vivo* MCMV infection

For *in vivo* MCMV infection, 8 week old *Xbp1*^fl/fl^ and *Xbp1*^ΔIEC^ mice were infected intraperitoneally with either 10^5^ PFU MCMV or with vehicle. During the infection mice were fed a standard chow diet and had free access to food and water. Five days after infection mice were sacrificed and the intestines were harvested.

### *In vitro* MCMV infection and quantification of replication

MCMV was prepared as described previously (*91*) and infection of Mode-K cells was performed at a MOI of 10. Fold induction of MCMV was quantified by qRT-PCR using the MCMV M54 or glycoprotein B (gB) gene sequence. M54: ATCATCCGTTGCATCTCGTTG (forward), CGCCATCTGTATCCGTCCAT (reverse) and gB: CTGGGCGAGAACAACGAGAT (forward), CGCAGCTCTCCCTTCGAGTA (reverse).

### Histology

After sacrifice, the small intestine was removed and immediately fixed in 10 % (w/v) formalin for 24h at 4°C. The tissue was then immersed in a series of ethanol solutions of increasing concentrations until 100 %. Next, the ethanol was gradually replaced with xylene, which is then replaced by paraffin. Paraffin embedded tissue was cut into 3.5-4.5 µm thin sections using the RM2255 microtome. Paraffin embedded tissue sections were stained for 2-5 min in haematoxylin. The cytoplasm was counterstained with 1 % (v/v) eosin solution for 2 min. Stained slides were then dehydrated and embedded in Roti-Histo-Kit mounting medium and examined with a Zeiss AxioImager.Z1 apotome microscope and the AxioVision Rel 4.9 software.

### Histology score CMV colitis

After staining, an expert pathologist assessed inflammation with a semi quantitative scoring system as previously reported (*3*). To assess the inflammation five subscores are summed up for every sample and multiplied by the extent of the affected area (1, <10%; 2, 10%-25%; 3, 25%-50%; 4, >50% of the intestine affected). As sub-scores the following parameters are used: mononuclear infiltrate (0: absent; 1: mild, 2: moderate 3: severe), crypt hyperplasia, epithelial injury/erosion, polymorphonuclear infiltrate and transmural inflammation.

### *Listeria monocytogenes* infection

*Listeria monocytogenes* serotype 1/2a strain EGD was used to test type I IFN induction in Mode-K cells. Bacteria were plated out from glycerol stocks onto brain-heart infusion (BHI; BD Biosciences, Franklin Lakes, USA) agar plates overnight. *Listeria* colonies were then transferred into 1 ml BHI medium and incubated overnight at 37°C to allow for bacterial replication. Next, bacteria were diluted and reincubated at 37°C to induce a log-phase growth until OD600 = 0.6 was reached. After washing with PBS, bacteria were diluted in Mode-K medium and added to the cells according to multiciply of infection (MOI). As described previously (*92*), Mode-K cells were infected at a MOI of 50. After allowing bacterial invasion for 1h, the bacteria-containing medium was removed, cells were washed once with PBS and then treated with gentamicin (50 µg/ml) for 1h to kill extracellular bacteria. The gentamicin-containing medium was then replaced with medium containing a lower concentration of Gentamicin (5 µg/ml), and cells were lysed for protein and RNA harvesting after another 22h.

### STING immunohistochemistry and quantification

To quantify STING expression in the intestinal epithelium, 3,3’-Diaminobenizidine staining was performed. Slides were rehydrated and boiled for 20 min in citrate buffer (pH 6.0) for antigen retrieval. Unspecific binding sites were blocked by incubating slides with 5 % (v/v) goat serum in phosphate-buffered saline (PBS). Naturally occurring peroxidases were inactivated by applying 3 % hydrogen peroxide. Antibody binding and 3,3’-diaminobenzidine staining was performed according to the manufacturer’s instruction (Vectastain Elite ABC Kit, Vector Labs, Peterborough, United Kingdom). In brief, after primary antibody binding, the signal was amplified by adding a horseradish peroxidase-conjugated streptavidin antibody. Quantification was performed by counting average STING positive cells per crypt. For each animal, 50 crypts were counted and averaged.

### Cell growth analysis

Cell growth was determined as measurement of the cell density (confluence) using the IncuCyte® Live-Cell Analysis system (Essen Bioscience) and following the manufacturer’s instructions. In brief, 5−10^4^ cells were seeded in 96 well plates as technical replicates (N = 6) and cultivated in DMEM for the indicated time period. After cell attachment, images were generated every 2h and the cell density was analysed using the IncuCyte® analysis software.

### Determination of GSH/GSSG ratio

GSH/GSSG ratios were obtained via a GSH/GSSG-GLO^TM^ assay kit (Promega, V6611) following the manufacturer’s instructions. For the experiments, 4−10^4^ iCtrl or *iXbp1* cells were plated out in opaque, white 96 well plates (Greiner, CELLSTAR, 655083), incubated for 24h in DMEM (10% FBS) and processed according to the manufacturès protocol. GSH/GSSG ratios were determine by the following equation: total GSH-(GSSGx2)/GSSG. All measurements were performed as technical replicates (N = 3) and repeated as independent, biological experiments (n = 3) indicated in the respective figure legend.

### Transcriptome data analysis of IBD bulk and scRNA sequencing datasets and definition of gene signatures as well as scores used in the study

The publicly available datasets GSE109142 (*46*) (RNA sequencing TPM counts of pediatric ulcerative colitis patients) and GSE73661 (*47*) (microarray of UC patients treated with either infliximab or vedolizumab) were downloaded from the Gene Expression Omnibus (GEO) repository and processed using *R* version 4.2.2. TPM counts were log_2_ transformed after adding a pseudo-count of 1 to remove zero values. If multiple microarray probe sets mapped to a same gene, the one exhibiting the highest variance was selected.

Digital gene expression matrices for SCP259 (*50*) (colon mucosa of 18 UC patients and 12 healthy individuals) as well as the metadata file containing the annotated cell barcodes were downloaded from the Single Cell Portal and processed in *R* using the Seurat package.

Gene expression scores were calculated using the R package singscore (v1.18.0) (*93, 94*) for the bulk datasets or as the mean of scaled log-normalised counts for the scRNA sequencing dataset. The ER-stress signature was calculated on 62 ER-stress related genes, as reported by Powell et al. (*12*). SSP score was calculated on the serine *de novo* synthesis pathway related genes *PHGDH*, *PSAT1* and *PSPH*. The cytoplasmic 1C cycle (CYTO) score was calculated on *MTHFD1*, *SHMT1* and *ALDH1L1* whereas *SHMT2, MTHFD2 and ALDH1L2 was used for* the mitochondrial 1C cycle (MITO) score.

The differential gene expression analysis performed on a monolayer of microdissected epithelial cells isolated from 5 healthy control and 7 UC patients was downloaded from the editor’s website (*48*).

### Stable isotope tracing and metabolite extraction

Stable isotope tracing experiments were performed as previously described (*44*). Briefly, [U-^13^C]-glucose or [U-^13^C]-glutamine (Cambridge Isotope Laboratories, CLM-1396 and CLM-1822) tracing was performed in DMEM supplemented with 10 % FBS and the respective tracers (2 mM glutamine or 17 mM glucose). 2−10^5^ cells were seeded in 12-well plates in triplicates for each experimental condition. Furthermore, an identical set of triplicate wells were plate-out to calculate the packed cell volume (PCV) by determining cell count and cell volume at the start and end of each tracing experiment (*36*). After 24h, growth medium was replaced by ^13^C tracer medium and cells were cultured for an additional 24h at 37°C. In parallel, starting PCV was determined by counting three wells per condition. To assess PCV at the end of the experiment, one set of triplicate wells was counted at the experimental endpoint, whereas another triplicates set was used for subsequent metabolite extraction. In addition, medium was collected, centrifuged at 350 g for 10 min at 4°C and supernatants were stored at −80°C until medium quantification via GC-MS or YSI.

For GC-MS analysis, cells from [U-^13^C]-glucose and [U-^13^C]-glutamine tracer experiments were washed once with 0.9 % ice-cold NaCl. Thereafter, metabolites were extracted by adding 400 μl ice-cold MeOH/mqH_2_O (ratio, 1:1) to each well. In addition, the extraction solvent contains the internal standards pentanedioc-d6 acid and [U-^13^C]-ribitol at a final concentration of 1 μg/ml as well as Tridecanoid-d25 acid at a final concentration of 5 μg/ml. After the plates were incubated for 5 min at 4°C on a rocking shaker, supernatant was transferred to 1.5-ml Eppendorf tubes, mixed with 200 μl CHCl_3_, and centrifuged for 5 min at 13,000 g and at 4°C. Finally, the upper polar phase was collected, fully dried via SpeedVac and stored at −20°C for subsequent GC-MS analysis. For LC-MS analysis, metabolites extraction was performed as previously described (*36*) and extracted metabolites were stored at −80°C until LC-MS analysis.

### GC-MS-based determination of MIDs of intracellular metabolites

Polar metabolites were derivatised for 90 min at 55°C with 20 μl of methoxyamine (c = 20 mg/ml) in pyridine under continuous shaking and subsequently for 60 min at 55°C with 20 μl of MTBSTFA w/ 1% TBDMCS. GC-MS analysis was performed using an Agilent 7890B GC coupled to an Agilent 5977A Mass Selective Detector (Agilent Technologies). A sample volume of 1 μl was injected into a Split/Splitless inlet, operating in splitless mode at 270°C. Gas chromatograph was equipped with a 30 m (I.D. 250 μm, film 0.25 μm) ZB-35MS capillary column with 5 m guard column (Phenomenex). Helium was used as carrier gas with a constant flow rate of 1.2 ml/min. GC oven temperature was held at 100°C for 2 min and increased to 300°C at 10°C/min and held for 4 min. Total run time was 26 min. Transfer line temperature was set to 280°C. Mass selective detector (MSD) was operating under electron ionisation at 70 eV. MS source was held at 230°C and the quadrupole at 150°C. For precise quantification of the MID, measurements were performed in selected ion monitoring mode. Target ions (m/z) and Dwell times are shown in Table S1.

The MetaboliteDetector software package (v. 3.220180913) was used for mass spectrometric data post processing, quantification, MID calculations, correction of natural isotope abundance, and determinations of fractional carbon contributions (*95*).

### Analysis of Medium Exchange Rates

Medium quantification and exchange rate calculation was performed as described in (*36*). For metabolite extraction, 20 µl of the collected medium sample was mixed with 180 µl extraction solvent (80 % MeOH/20 % mqH_2_O) containing the internal standards [U-13C]-ribitol (50 µg/ml) and pentanedioc-d6 acid (20 µg/ml), vortexed for 10 s and subsequently incubated for 15 min at 4°C and 1350 rpm in a thermomixer. After centrifugation for 5 min at max speed, 50 µl of the supernatant was transferred in GC-vial and dried on SpeedVac overnight.

Metabolite derivatisation was performed by using a multi-purpose sample preparation robot (Gerstel). Dried medium extracts were dissolved in 30 µl pyridine, containing 20 mg/mL methoxyamine hydrochloride (Sigma-Aldrich), for 120 min at 45°C under shaking. After adding 30 µl N-methyl-N-trimethylsilyl-trifluoroacetamide (Macherey-Nagel), samples were incubated for 30 min at 45°C under continuous shaking.

GC-MS analysis was performed by using an Agilent 7890A GC coupled to an Agilent 5975C inert XL Mass Selective Detector (Agilent Technologies). A sample volume of 1 µl was injected into a Split/Splitless inlet, operating in split mode (10:1) at 270°C. The gas chromatograph was equipped with a 30 m (I.D. 0.25 mm, film 0.25 µm) DB-5ms capillary column (Agilent J&W GC Column) with 5 m guard column in front of the analytical column. Helium was used as carrier gas with a constant flow rate of 1.2 ml/min. The GC oven temperature was held at 90°C for 1 min and increased to 220°C at 10°C/min. Then, the temperature was increased to 280°C at 20°C/min followed by 5 min post run time at 325°C. The total run time was 22 min. The transfer line temperature was set to 280°C. The MSD was operating under electron ionisation at 70 eV. The MS source was held at 230°C and the quadrupole at 150°C. Mass spectra were acquired in selected ion monitoring (SIM) mode for precise quantification of medium components. In Table S2, masses used for quantification and qualification of the derivatised target analytes (dwell times between 25 and 70 ms) are illustrated.

All GC-MS chromatograms were processed using MetaboliteDetector (v. 3.2.20190704). Compounds were annotated by retention time and mass spectrum using an in-house mass spectral (SIM) library (overall similarity: >0.60). The following deconvolution settings were applied: Peak threshold: 2; Minimum peak height: 2; Bins per scan: 10; Deconvolution width: 8 scans; no baseline adjustment; Minimum 1 peaks per spectrum; no minimum required base peak intensity.

Peak areas of all isotopologues of defined quantification ions were summed up and divided by the peak area of the internal standard for normalisation. In addition, a calibration curve was prepared to calculate absolute concentrations. Absolute uptake and release rates were calculated as described previously (*95*).

### LC-MS measurements

Untargeted LC-MS analysis was carried out as previously (*36*). The method was used to detect glutathione, cystine and γ-L-glutamyl-L-cysteine in cell extracts. For calculating intracellular metabolite level, measured peak areas were initially corrected for natural isotope abundance, normalised to internal standards and subsequently divided by PCV to obtain relative metabolite levels.

### YSI measurements

Collected medium samples were initially filtrated (Phenex-RC 4mm Syringe Filters 0.2 µm) prior measurement to remove particles. Absolute quantitative values for glucose, lactate, glutamine and glutamic acid were acquired using the YSI 2950D Biochemistry Analyzer (Kreienbaum KWM). For a precise and reliable quantification, external calibration curves of each compound were prepared and measured in duplicates. To get absolute consumption and release rate (CORE, fmol/PCV/h), measured concentrations were divided by PCV (determined at start and end of the experiment, see stable isotope tracing for details) and the duration of the experiment.

### Flow Cytometric Analysis of LPO Levels

Image-iT™ Lipid Peroxidation Kit (ThermoFisher) was used according to manufacturer’s instructions. In brief, 250,000 iCtrl or *iXbp1* Mode-K cells were cultivated in 2 ml medium and treated after 24h as stated in the figure legends. Following 24h incubation, cells were harvest via trypsinization, centrifuged at 350 g for 5 min and washed with warm DMEM. Then, cells were stained in 100 μl DMEM supplemented with 10 µM Lipid peroxidation sensor (BODIPY® 581/591 C11) for 30 min at 37°C. After two washing steps with ice-cold PBS at 350 g for 5 min, cells were resuspended in ice-cold 400 µl FACS buffer and measured using NovoCyte Quanteon system (Agilent). Data analysis was performed using FlowJo v10.6.2 software. All measurements were performed as independent, biological experiments (n ≥ 3).

### Flow Cytometric Analysis of cytoplasmic ROS Levels

250,000 iCtrl or *iXbp1* Mode-K cells were seeded in six-well plates and treated the subsequent day as indicated in the figure legends. After an incubation time of 24h, cells were stained with Zombie NIR (BioLegend, 1:1000 diluted in PBS) for 20 min at 37°C (to discriminate live and dead cells in the later analysis), detached with trypsin, centrifuged at 350 g for 5 min and washed with warm DMEM. Then, cells were stained in 100 μl DMEM supplemented with 10 µM DCF-DA (PromoCell) for 30 min at 37°C. After the incubation step, cells were centrifuged at 350 g for 5 min, washed twice with ice-cold PBS and resuspended in ice-cold FACS buffer (PBS with 10% FBS and 0.5 mM EDTA). Samples were measured using the BD FACSCanto^TM^ system and BD FACSDiva software. Data analysis was performed using FlowJo v10.6.2 software. All measurements were performed as independent, biological experiments (n ≥ 3) indicated in the respective figure legend.

### Western blot analysis

Cells and tissues were lysed using 40-100 μl SDS-based DLB buffer + 1% Halt™ Protease inhibitor cocktail (ThermoFisher Scientific, Waltham, USA) before incubation for 5 min at 95°C followed by sonification for 2x 5 s. To remove cell remnants, lysates were centrifuged at 16.000 x g for 15 min at 4°C and supernatants were transferred into a new tube. For protein extraction of organoids, Matrigel™ was removed by incubation of organoids in Corning™ Cell Recovery Solution (Corning, New York, USA) for 45-60 min on ice after transferring organoids to tubes, centrifugation at 3000 g for 2 min and discarding supernatants. Next, the lysis buffer as described above was added. The protein concentration was determined applying the copper-based DC Protein Assay (Bio-Rad, Munich, Germany) according to the manufactureŕs protocol. Proteins were separated by their molecular weight using precast gradient polyacrylamide gels for improved separation of a wide range of proteins (NuPAGE® 4-12% Bis-Tris gels, Life Technologies, Darmstadt, Germany). Protein lysates were prepared by supplementation with the corresponding NuPAGE LDS Sample Buffer (Thermo Scientific, Bremen, Germany), followed by incubation at 70°C for 70 min. SDS-PAGE electrophoresis was run at 160 V and a maximum current of 300 mA for 1 to 2h in the XCell SureLock Mini Cell system (Life Technologies, Darmstadt, Germany). NuPAGE MOPS SDS running buffer was used to run the gels and Prestained PageRuler plus protein ladder 10-250 K (Thermo Scientific, Bremen, Germany) served to estimate protein size. To detect proteins using immunodetection by specific antibodies, separated proteins were transferred onto polyvinylidene difluoride (PVDF) membranes. PVDF membranes (Bio-Rad, Munich, Germany) were activated under mild agitation using methanol for 10 s, followed by 5 min of washing in mqH_2_O and then transferred for 5 min to anode buffer 1. Separated proteins were transferred from the gel onto the membrane by semi-dry blotting using the Trans-Blot Turbo™ Transfer System (Bio-Rad, Munich, Germany) with a discontinuous buffer system consisting of one cathode buffer and two anode buffers. Proteins were blotted onto the membrane at 0,3 A, 25 V for 60 min. After the transfer membranes were blocked with 5% (w/v) blotting grade blocker (non-fat dry milk) in Tris - buffered saline (TBS) supplemented with 0.1% (v/v) Tween 20 (TBS - T) for 1h. Membranes were probed with the appropriate primary and secondary antibody. The primary antibody was applied in 5% (w/v) blotting grade blocker (non-fat dry milk) or bovine serum albumin (BSA) in TBS-T according to the manufacturer’s instruction and incubated overnight at 4°C under mild shaking. Next, the membrane was washed three times with TBS - T for 15 min to remove excess antibody. A horseradish peroxidase (HRP)-or IRDye-conjugated secondary antibody was diluted in 5% (w/v) blotting grade blocker (non-fat dry milk) in TBS-T according to the manufacturer’s recommendation and incubated for 1h at room temperature. Binding of the secondary HRP-coupled antibody was detected using a chemiluminescent substrate kit (Thermo Scientific) and recorded with an automated developer machine (Agfa). Detection of IRDye-conjugated antibodies were recorded with an Odyssey imager system (Li-Cor Biosciences). The following antibodies were used: Actin β (Sigma-Aldrich, A-5441), β-Tubulin (MBL International, M150-3), Vinculin (Cell Signaling Technology, 13901), Aldh1l2 (Sigma-Aldrich, HPA039481), phospho-Sting (Cell Signaling Technology, 72971), phospho-Tbk1 (Cell Signaling Technology, 5483), Sting (Cell Signaling Technology, 13647), Tbk1 (Cell Signaling Technology, 38066), mouse-HRP (Amersham Biosciences, NA931V), rabbit-HRP (Amersham Biosciences, NA934V), rabbit-IRDye® 800CW (Li-COR Technologies, 926-32213).

### Quantitative real-time PCR (qRT-PCR) analysis

RNA extraction was performed using the RNeasy Mini Kit (Qiagen, Hilden, Germany) according to the manufacturer’s instruction. To generate cDNA, 100-1000 ng total RNA (depending on the amount of previously extracted RNA) was transcribed into complementary DNA (cDNA) by reverse transcription using the Maxima H Minus First Strand cDNA Synthesis Kit (Thermo Scientific, Bremen, Germany). Reverse transcription was performed following the manufactureŕs instruction. qRT-PCR was used to amplify and simultaneously quantify mRNA levels using target-specific oligonucleotides. Predesigned TaqMan probes (Applied Biosystems, Carlsbad, USA were purchased and used. The 7900HT Fast Real Time PCR System (Applied Biosystems, Darmstadt, Germany) was used for qRT-PCR experiments. Samples were run in duplicates on 384-well plates. For the PCR reaction, 5-10 ng cDNA and 0.5 μl of the respective TaqMan gene expression assays were used. The PCR program was carried out following the manufacturer’s recommendation (TaqMan Gene Expression Master Mix protocol, Applied Biosystems, Darmstadt, Germany).

Alternatively, qRT-PCR was performed from 10 ng cDNA per sample using Fast SYBR™ Green Master Mix (Lifetech) at 95°C for 20 s, 40 cycles of 95°C for 1 s and 60°C for 20 s on the QuantStudio 5 Real-Time PCR System (ThermoFisher). Relative quantification of each gene was done by QuantStudio Design&Analysis v1.5.1 software (ThermoFisher) and comparative Ct method. All samples were analysed in technical duplicates and repeated as independent, biological experiments (n ≥ 3). Moreover, all expression data were normalised to housekeeping genes (*Tbp*, *Hrpt1* or *Actinb*). Taqman assay probes were purchased from Life Technologies (*Cxcl10*, 04331182; *Gapdh*, 99999915; Ifnb1, 04331182; *Tmem173*, 04331182; *Xbp1*, 03464496) or Integrated DNA Technologies (*Shmt1* NM_009171(1)*; Shmt2* NM_001252316(2); *Aldh1l1* NM_027406(1); *Aldh1l2* NM_153543(1); *Mthfd1* NM_138745(1); *Mthfd2* NM_008638(1); *Psat1* NM_001205339(2); *Psph* NM_133900(1); *Phghd* NM_016966(1) and *Actb* NM_007393(1)). All non-TaqMan assay mouse primers used in this study can be found in the Table S3:

### Quantification and statistical analysis

GraphPad Prism 5 software package (GraphPad Software Inc., La Jolla, USA) was used for statistical analysis. If not otherwise specified, an unpaired, two-tailored Student’s t-test was performed. Data are visualised as mean ± standard error of the mean (SEM), if not otherwise stated. *P* values of <0.05 (*), <0.01 (**) or <0.001 (***) were considered statistically significant.

We define one *n* as one independent biological experiment (in some cases further consisting of several wells, e.g. triplicate wells for all stable isotope tracing experiments). The technical mean of one biological experiment was considered as one *n*. Mean values of several independent, biological experiments (as indicated in figure legends) were plotted and used for statistical analysis as indicated.

## Acknowledgments

We thank M. Rohm, M. Hansen, S. Kock, D. Oelsner, S. Baumgarten, M. Reffelmann, J. Ohrndorf for perfect technical assistance. We also thank the LCSB Metabolomics Platform, especially Xiangyi Dong and Floriane Vanhalle, for providing technical and analytical support; the National Cytometry Platform (Quantitative Biology Unit, LIH) and especially Thomas Cerutti for support with flow cytometric analyses.

## Author contributions

PR, KA, JM, FW and BB designed the study. BB, FW, MB, LM, JK, GY, SJW, LN, SB, NK, FT, LW, JW, ZB, STS, GI, OIC, CK, CJ, EK, EL, DH, SRP, TA, JM, and KA performed experiments and analysed data. LCS, RB, AK and SSc provided intellectual input and guidance. PR, KA, JM, FW and BB planned the project and supervised the experiment. PR, KA, JM, FW and BB wrote the manuscript.

## Data and materials availability

All data needed to evaluate the conclusions in the paper are present in the paper and/or the Supplementary Materials.

## Supplementary Materials

**Fig. S1:**
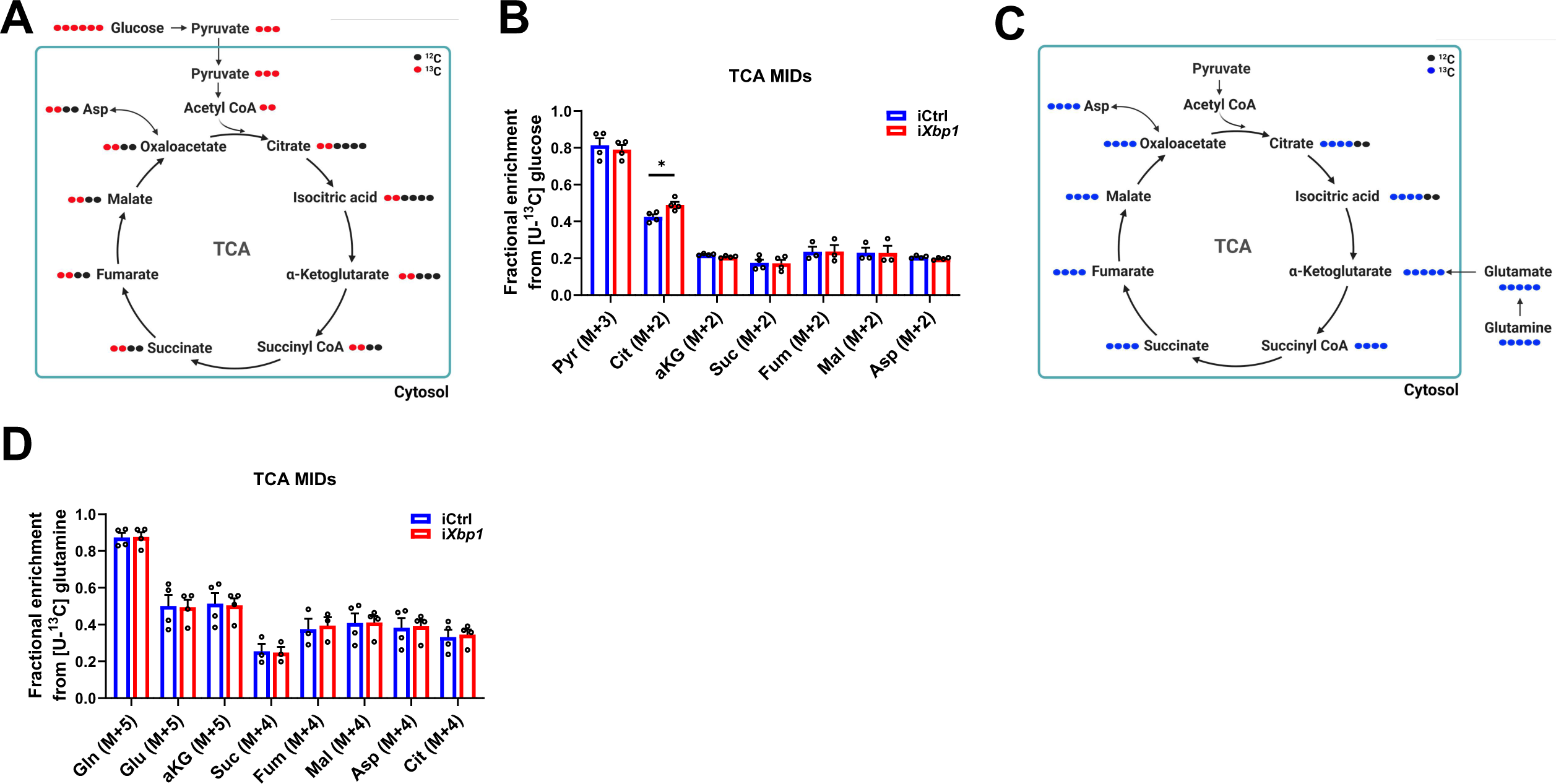
IECs show a metabolic reprogramming towards GSH metabolism to cope with ER stress. (**A**) Schematic illustration of the mass isotopomer distribution (MID) of the TCA cycle from glucose. Filled red circles represent ^13^C atoms derived from [U-^13^C]-glucose. (**B**) Relative MIDs of the indicated TCA cycle intermediates in iCtrl or i*Xbp1* cells cultured in [U-^13^C]-glucose medium for 24h (n ≥ 3). (**C**) Schematic illustration of the mass isotopomer distribution (MID) of the TCA cycle from glutamine. Filled blue circles represent ^13^C atoms derived from [U-^13^C]-glutamine. (**D**) Relative MIDs of the indicated TCA cycle intermediates in iCtrl or i*Xbp1* cells cultured in [U-^13^C]-glutamine medium for 24h (n ≥ 3). \**P*<0.05, \*\**P* <0.01, \*\*\**P* <0.001.

**Fig. S2:**
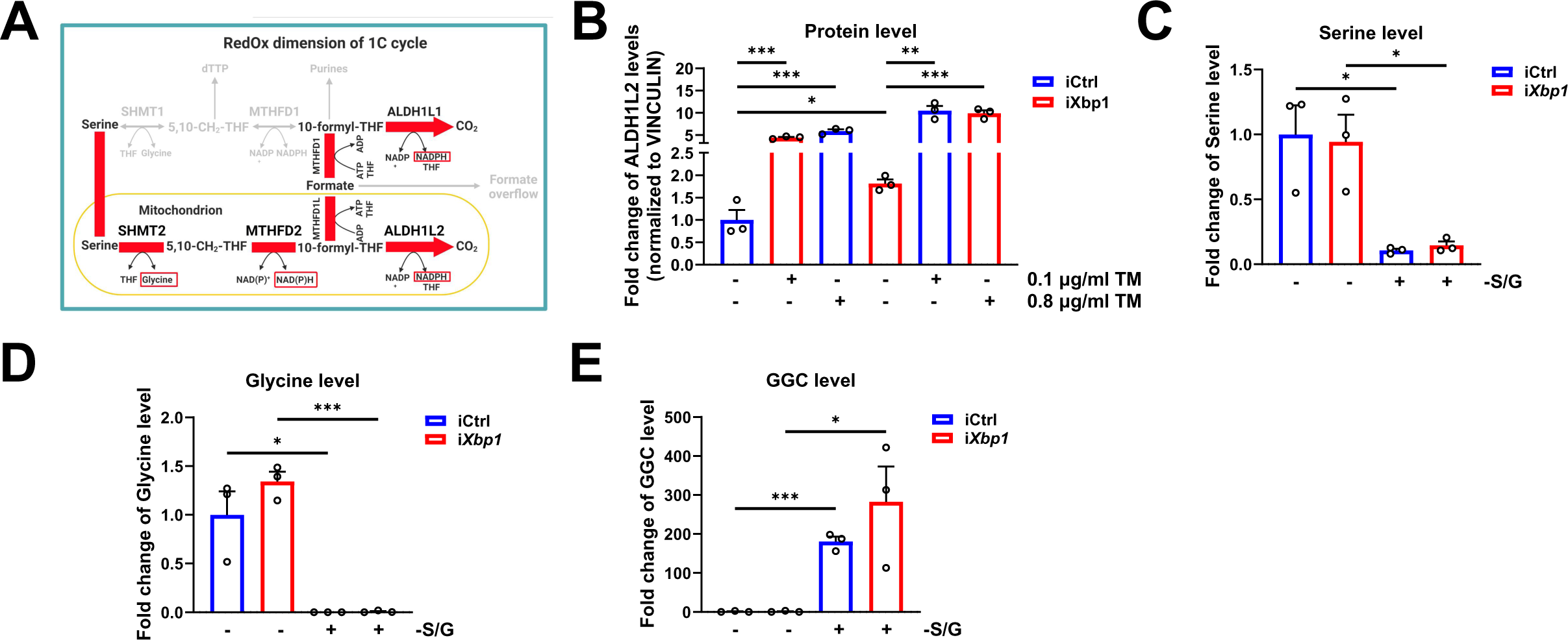
Chronic ER-stress polarizes the 1C metabolism towards glycine and NADPH supply. (**A**) Schematic model of the theoretical contribution of serine to the cellular RedOx homeostasis via the 1C cycle under ER-stress conditions. (**B**) Quantification of ALDH1L2 expression in TM-treated iCtrl and i*Xbp1* cells (n = 3, independent Western blots). ALDH1L2 expression was normalized to the loading control VINCULIN. (**C-E**) Intracellular metabolite level of (C) serine, (D) glycine and (E) γ-glutamyl-cysteine (GGC) in iCtrl and i*Xbp1* cells cultured in complete or serine/glycine-free (-S/G) medium for 24 h (n=3). \**P*<0.05, \*\**P* <0.01, \*\*\**P* <0.001.

**Fig.S3:**
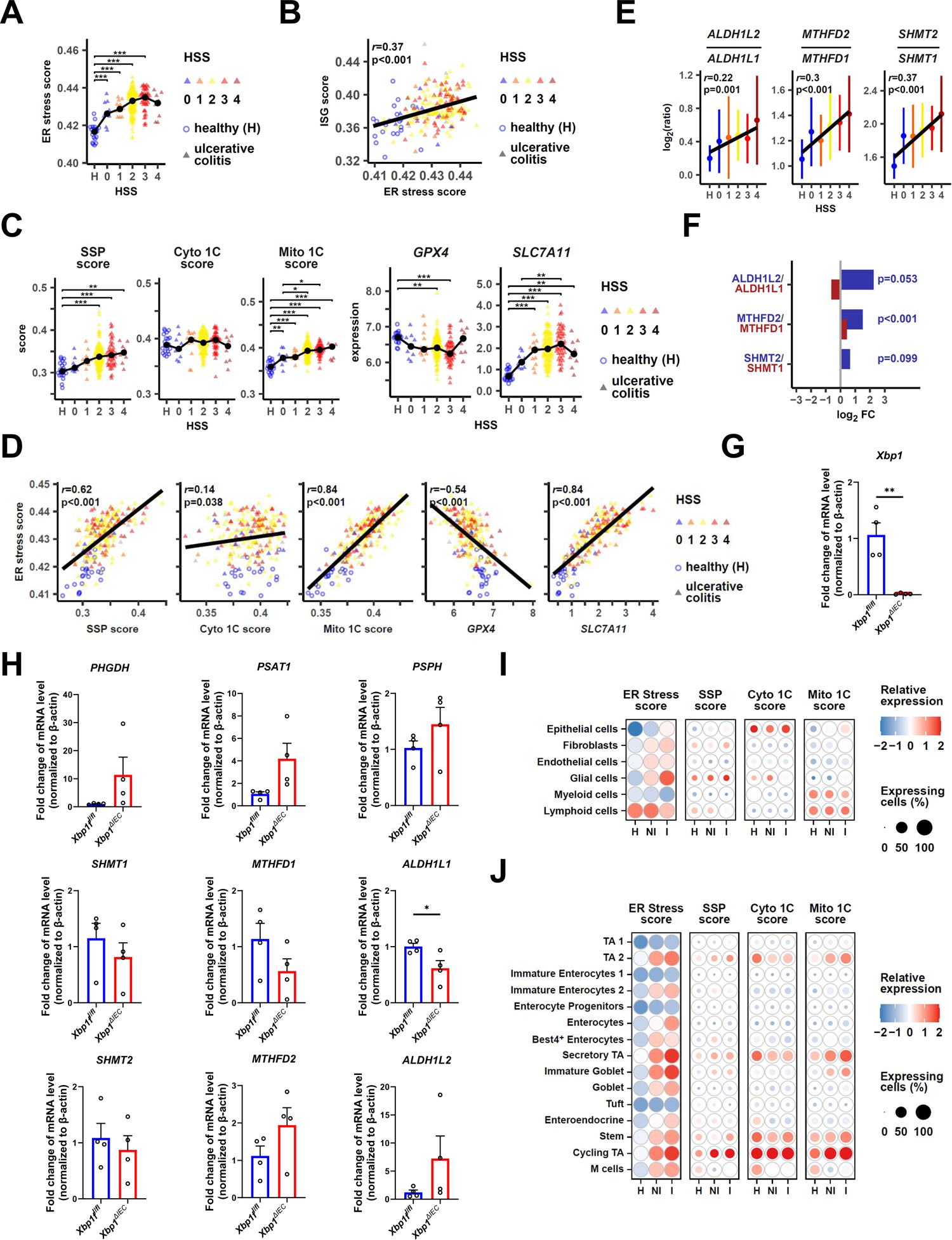
Metabolic rewiring of the SGOC metabolism is taking place in the inflamed mucosa of IBD patients. (**A**) Correlation of ER-stress with histology severity scores (HSS) in UC patients using the GSE109142 dataset. Significances of the Tukey HSD post-hoc test following a one-way ANOVA are shown. (**B**) Correlation of ER-stress with the interferon-stimulating gene signature in the GSE109142 dataset. Pearson’s correlation coefficients *r* and p-values are shown. **(C**) Expression and scores of SGOC related genes in UC patients from the GSE109142 dataset. Black dots and connecting lines show the median expression. Significances of the Tukey HSD post-hoc test following a one-way ANOVA are shown. (**D**) Correlation of the indicated scores and genes with ER-stress and disease severity in the GSE109142 dataset. Pearson’s correlation coefficients *r* and p-values are shown. (**E**) Gene expression ratios of the mitochondrial 1C cycle enzymes over their cytosolic counterparts (*ALDH1L2*/*ALDH1L1*, *SHMT2/SHMT1*, *MTHFD2*/*MTHFD1* in the GSE109142 dataset and their correlation with IBD severity. (**F**) Log_2_ fold changes of mRNA expression of the mitochondrial and cytosolic 1C cycle gene pairs in colon biopsies of UC patients compared to healthy controls. (**G**) Relative *Xbp1* expression level in small-intestinal crypts derived from *Xbp1*^fl/fl^ and *Xbp1*^ΔIEC^ mice (n = 4). (**H**) Relative mRNA expression level of the indicated genes related to SSP and the mitochondrial (mito 1C) and cytoplasmic (cyto 1C) 1C cycle in small-intestinal crypts derived from *Xbp1*^fl/fl^ and *Xbp1*^ΔIEC^ mice (n = 4). (**I**) Heatmap of the indicated scores (ER-stress, SSP, mito 1C and Cyto 1C) in the main cell-types of the SCP259 scRNA-Seq IBD dataset in healthy, non-inflamed (NI) and inflamed (I) biopsies (colon mucosa of 18 UC patients and 12 healthy individuals (*1*)). Colour shows z-scores and the dot size the percent of expressing cells. (**J**) Heatmap of the indicated scores in epithelial cell clusters of SCP259 in healthy, non-inflamed (NI) and inflamed (I) colon biopsies. Colour shows z-scores and the dot size the percent of expressing cells. *p<0.05, **p<0.01, ***p<0.001.

**Fig. S4:**
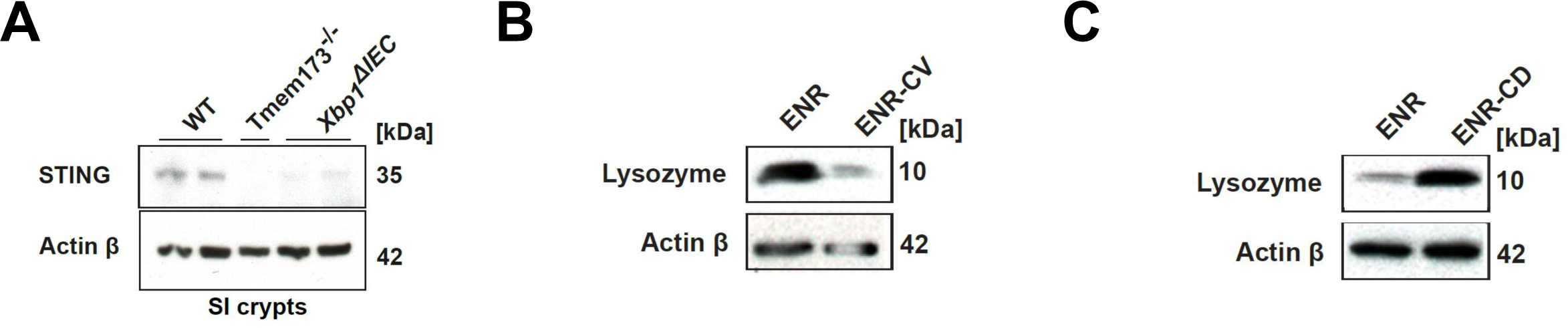
Chronic ER-stress impairs cGAS/STING signalling *in vitro* and *in vivo*. (**A**) STING expression in SI crypts isolated from *Xbp1*^fl/fl^ (n=2), *Tmem173* −/− (n=1) and *Xbp1*^ΔIEC^ (n=2) mice. (**B+C**) Western blot analysis of lysozyme expression in SI organoids enriched for stem cells (B) or paneth cells (C) compared to enterocytes-enriched organoid cultures.

**Fig. S5:**
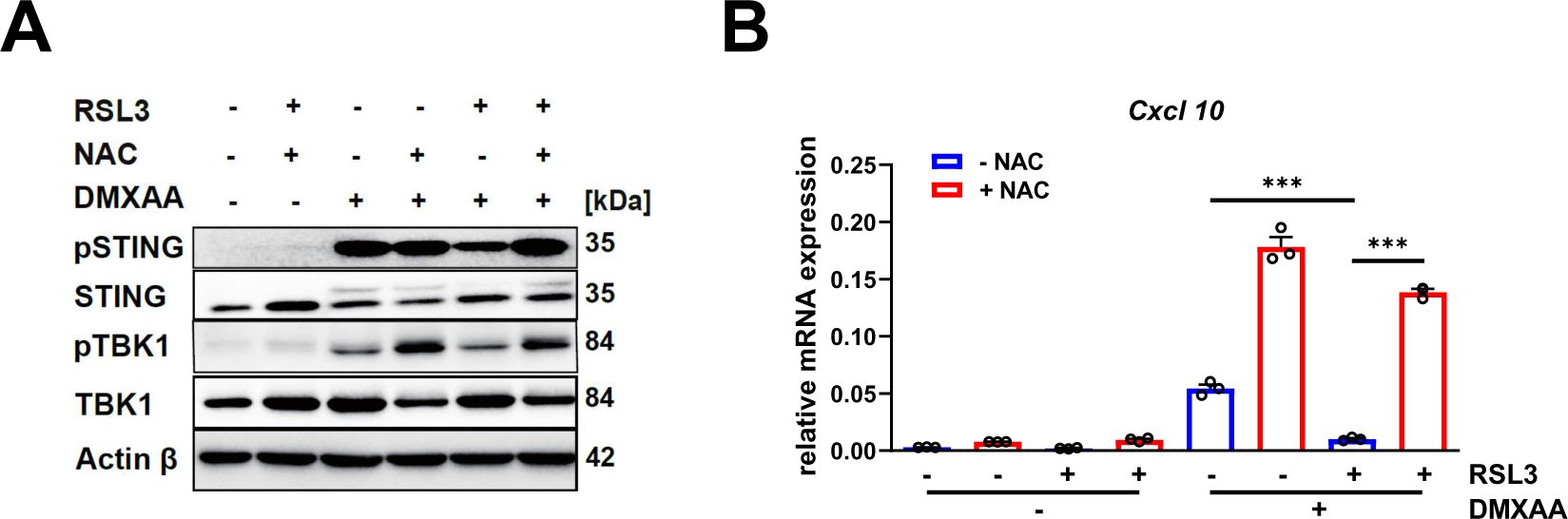
RSL3-induced cGAS/STING inhibition is rescued by antioxidant treatment. (**A**) Western Blot analysis of iCtrl cells stimulated with DMXAA (100 μg/ml, 1 h) after 24 h pretreatment with 10 mM NAC and 0.5 μM RSL3. (**B**) *Cxcl10* expression of iCtrl cells stimulated with DMXAA (100 µg/ml, 3 h) after 24 h of co-treatment with RSL3 (0.5 μM) and NAC (10 mM) (n=3). \**P*<0.05, \*\**P* <0.01, \*\*\**P* <0.001.

**Table S1:** Target ions and Dwell times of metabolites quantified in this study.

| **Derivative**<br>**(m/z of selected fragment)** | **Quant-Ions**<br>**(m/z)** | **Dwell Time (ms/**<br>**for each)** | **Sum Formula**<br>**(fragment)** | **MID**<br>**size** |
| --- | --- | --- | --- | --- |
| **Pyruvic Acid 1MeOX 1TBDMS (174)** | 174.0 – 180.1 | 15 | C6H12O3NSi | 4 |
| **Lactic acid 2TBDMS (261)** | 261.1 – 267.1 | 15 | C11H25O3Si2 | 4 |
| **Alanine 2TBDMS (260)** | 260.1 – 266.1 | 15 | C11H26NO2Si2 | 4 |
| **Glycine 2TBDMS (246)** | 246.1 – 252.1 | 15 | C10H24NO2Si2 | 3 |
| **Valine 2TBDMS (288)** | 288.2 – 295.2 | 15 | C13H30NO2Si2 | 6 |
| **Leucine 2TBDMS (302)** | 302.2 – 310.2 | 15 | C14H32NO2Si2 | 7 |
| **Isoleucine 2TBDMS (302)** | 302.2 – 310.2 | 15 | C14H32NO2Si2 | 7 |
| **Proline 2TBDMS (286)** | 286.2 – 294.2 | 15 | C13H28NO2Si2 | 6 |
| **Succinic Acid 2TBDMS (289)** | 289.1 – 296.1 | 15 | C12H25O4Si2 | 5 |
| **Fumaric Acid 2TBDMS (287)** | 287.1 – 294.1 | 15 | C12H23O4Si2 | 5 |
| **(Internal Standard) Pentanedioic-D6**<br>**Acid** | 235.2, 309.2,<br>351.3 | 30 |  |  |
| **Serine 3TBDMS (390)** | 390.2 – 396.2 | 15 | C17H40NO3Si3 | 4 |
| **Methionine 2TBDMS (320)** | 320.2 – 328.2 | 10 | C13H30NO2SSi2 | 6 |
| **α-Ketoglutaric Acid 1MeOX**<br>**2TBDMS (346)** | 346.2 – 354.2 | 10 | C14H28NO5Si2 | 6 |
| **Malic Acid 2TBDMS (419)** | 419.2 – 426.2 | 15 | C18H39O5Si3 | 5 |
| **Aspartic Acid 3TBDMS (418)** | 418.2 – 425.2 | 15 | C18H40NO4Si3 | 5 |
| **Glutamic Acid 3TBDMS (432)** | 432.3 – 440.3 | 10 | C19H42NO4Si3 | 6 |
| **Lysine 3TBDMS (431)** | 431.3 – 440.3 | 15 | C20H47N2O2Si3 | 7 |
| **Glutamine 3TBDMS (431)** | 431.3 – 439.3 | 10 | C19H43N2O3Si3 | 6 |
| **Citric Acid 4TBDMS (591)** | 591.3 – 600.3 | 10 | C26H55O7Si4 | 7 |
| **3-Phosphoglyceric acid 4TBDMS**<br>**(585)** | 585.4 – 592.4 | 15 | C27H63O7PSi4 | 4 |
| **Histidine 3TBDMS (440)** | 440.2 – 449.2 | 15 | C20H42N3O2Si3 | 7 |
| **Tryptophan 2TBDMS (375)** | 375.2 – 389.2 | 10 | C19H31N2O2Si2 | 12 |

**Table S2:** List of quantification and qualification ions used for metabolite detection in medium.

| Analyte Name | Quantification Ions (*m/z*) | Qualification Ion I (*m/z*) | Qualification Ion II (*m/z*) |
| --- | --- | --- | --- |
| Lactic acid 2TMS | 219.1-224.1 | 190.1 | 117.1 |
| Alanine 2TMS | 218.1-223.1 | 190.1 | 116.1 |
| Leucine 2TMS | 158.1 | 218.1 | 232.2 |
| Isoleucine 2TMS | 158.1 | 218.1 | 232.2 |
| Glycine 3TMS | 276.1-280.1 | 248.1 | 174 |
| Serine 3TMS | 306.1-311.1 | 218.1 | 204.1 |
| Threonine 3TMS | 218.1 | 291.2 | 320.2 |
| IS Pentanedioic acid-D6 2TMS | 267.1 | 163.1 | 239.1 |
| Methionine 2TMS | 176.1 | 250.1 | 293.1 |
| Glutamic acid 3TMS | 363.2-371.2 | 246.1 | 348.2 |
| Phenylalanine 2TMS | 192.1 | 218.1 | 266.1 |
| IS [UL-13C5]-Ribitol 5TMS | 220.1 | 310.2 | 323.2 |
| Glutamine 3TMS | 347.2-354.2 | 245.1 | 156.1 |
| Glucose 1MEOX 5TMS | 319.2, 323.2 | 217.1 | 220.1 |
| Lysine 4TMS | 174.1 | 317.2 | 434.3 |
| Tyrosine 3TMS | 218.1 | 179.1 | 280.2 |
| Tryptophan 2TMS | 218.1-223.1 | 130 | 348.2 |
| Cystine 4TMS | 218.1-223.1 | 297.1 | 411.2 |

**Table S3:**
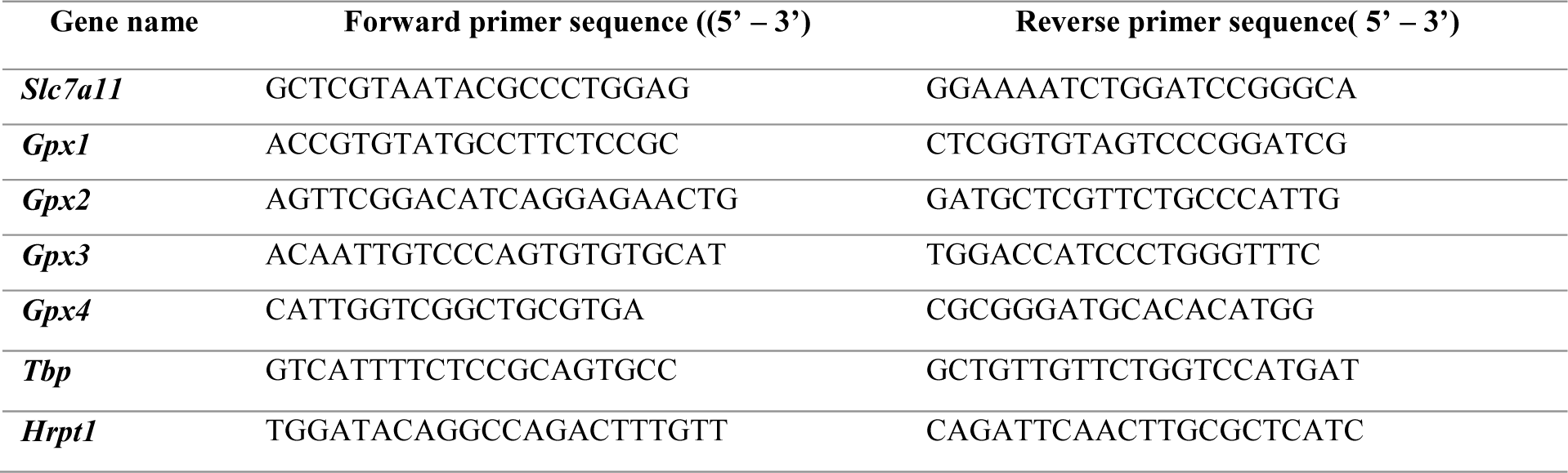
List of forward and reverse primer used for qRT-PCR analysis.

## References

1. G. G. Kaplan, The global burden of IBD: from 2015 to 2025. Nat Rev Gastroenterol Hepatol 12, 720–727 (2015).

2. J. Grootjans, A. Kaser, R. J. Kaufman, R. S. Blumberg, The unfolded protein response in immunity and inflammation. Nat Rev Immunol 16, 469–484 (2016).

3. T. E. Adolph et al., Paneth cells as a site of origin for intestinal inflammation. Nature 503, 272–276 (2013).

4. C. K. Heazlewood et al., Aberrant mucin assembly in mice causes endoplasmic reticulum stress and spontaneous inflammation resembling ulcerative colitis. PLoS Med 5, e54 (2008).

5. A. Kaser et al., XBP1 links ER stress to intestinal inflammation and confers genetic risk for human inflammatory bowel disease. Cell 134, 743–756 (2008).

6. F. Zhao et al., Disruption of Paneth and goblet cell homeostasis and increased endoplasmic reticulum stress in Agr2−/− mice. Dev Biol 338, 270–279 (2010).

7. C. Hetz, The unfolded protein response: controlling cell fate decisions under ER stress and beyond. Nat Rev Mol Cell Biol 13, 89–102 (2012).

8. D. Ron, P. Walter, Signal integration in the endoplasmic reticulum unfolded protein response. Nat Rev Mol Cell Biol 8, 519–529 (2007).

9. O. I. Coleman, D. Haller, ER Stress and the UPR in Shaping Intestinal Tissue Homeostasis and Immunity. Front Immunol 10, 2825 (2019).

10. S. P. Eugene, V. S. Reddy, J. Trinath, Endoplasmic Reticulum Stress and Intestinal Inflammation: A Perilous Union. Front Immunol 11, 543022 (2020).

11. S. S. Cao, Cellular Stress Responses and Gut Microbiota in Inflammatory Bowel Disease. Gastroenterol Res Pract 2018, 7192646 (2018).

12. N. Powell et al., Interleukin-22 orchestrates a pathological endoplasmic reticulum stress response transcriptional programme in colonic epithelial cells. Gut 69, 578–590 (2020).

13. W. Vanhove et al., Biopsy-derived Intestinal Epithelial Cell Cultures for Pathway-based Stratification of Patients With Inflammatory Bowel Disease. J Crohns Colitis 12, 178–187 (2018).

14. L. Welz et al., Epithelial X-Box Binding Protein 1 Coordinates Tumor Protein p53-Driven DNA Damage Responses and Suppression of Intestinal Carcinogenesis. Gastroenterology 162, 223–237 e211 (2022).

15. L. Niederreiter et al., ER stress transcription factor Xbp1 suppresses intestinal tumorigenesis and directs intestinal stem cells. J Exp Med 210, 2041–2056 (2013).

16. M. Benzarti, C. Delbrouck, L. Neises, N. Kiweler, J. Meiser, Metabolic Potential of Cancer Cells in Context of the Metastatic Cascade. Cells 9, (2020).

17. H. P. Harding et al., An integrated stress response regulates amino acid metabolism and resistance to oxidative stress. Mol Cell 11, 619–633 (2003).

18. H. Sies, D. P. Jones, Reactive oxygen species (ROS) as pleiotropic physiological signalling agents. Nat Rev Mol Cell Biol 21, 363–383 (2020).

19. Z. Li, D. Xu, X. Li, Y. Deng, C. Li, Redox Imbalance in Chronic Inflammatory Diseases. Biomed Res Int 2022, 9813486 (2022).

20. W. Luczaj, A. Gegotek, E. Skrzydlewska, Antioxidants and HNE in redox homeostasis. Free Radic Biol Med 111, 87–101 (2017).

21. M. Jia et al., Redox homeostasis maintained by GPX4 facilitates STING activation. Nat Immunol 21, 727–735 (2020).

22. M. Motwani, S. Pesiridis, K. A. Fitzgerald, DNA sensing by the cGAS-STING pathway in health and disease. Nat Rev Genet 20, 657–674 (2019).

23. K. P. Hopfner, V. Hornung, Molecular mechanisms and cellular functions of cGAS-STING signalling. Nat Rev Mol Cell Biol 21, 501–521 (2020).

24. F. Wottawa, D. Bordoni, N. Baran, P. Rosenstiel, K. Aden, The role of cGAS/STING in intestinal immunity. Eur J Immunol 51, 785–797 (2021).

25. C. W. Lio et al., cGAS-STING Signaling Regulates Initial Innate Control of Cytomegalovirus Infection. J Virol 90, 7789–7797 (2016).

26. S. J. Piersma et al., Virus infection is controlled by hematopoietic and stromal cell sensing of murine cytomegalovirus through STING. Elife 9, (2020).

27. M. Stempel et al., The herpesviral antagonist m152 reveals differential activation of STING-dependent IRF and NF-kappaB signaling and STING’s dual role during MCMV infection. EMBO J 38, (2019).

28. Y. Qin et al., Risk Factors of Cytomegalovirus Reactivation in Ulcerative Colitis Patients: A Meta-Analysis. Diagnostics (Basel) 11, (2021).

29. E. S. Guimaraes et al., Brucella abortus Cyclic Dinucleotides Trigger STING-Dependent Unfolded Protein Response That Favors Bacterial Replication. J Immunol 202, 2671–2681 (2019).

30. J. Moretti et al., STING Senses Microbial Viability to Orchestrate Stress-Mediated Autophagy of the Endoplasmic Reticulum. Cell 171, 809–823 e813 (2017).

31. K. Aden et al., ATG16L1 orchestrates interleukin-22 signaling in the intestinal epithelium via cGAS-STING. J Exp Med 215, 2868–2886 (2018).

32. X. Ma et al., Intestinal Epithelial Cell Endoplasmic Reticulum Stress and Inflammatory Bowel Disease Pathogenesis: An Update Review. Front Immunol 8, 1271 (2017).

33. A. Kaser, E. Martinez-Naves, R. S. Blumberg, Endoplasmic reticulum stress: implications for inflammatory bowel disease pathogenesis. Curr Opin Gastroenterol 26, 318–326 (2010).

34. W. Zheng et al., Evaluation of AGR2 and AGR3 as candidate genes for inflammatory bowel disease. Genes Immun 7, 11–18 (2006).

35. J. L. Parker et al., Molecular basis for redox control by the human cystine/glutamate antiporter system xc(). Nat Commun 12, 7147 (2021).

36. J. Meiser et al., Serine one-carbon catabolism with formate overflow. Sci Adv 2, e1601273 (2016).

37. C. F. Labuschagne, N. J. van den Broek, G. M. Mackay, K. H. Vousden, O. D. Maddocks, Serine, but not glycine, supports one-carbon metabolism and proliferation of cancer cells. Cell Rep 7, 1248–1258 (2014).

38. M. Benzarti et al., PKM2 diverts glycolytic flux in dependence on mitochondrial one-carbon cycle. bioRxiv, 2023.2001.2023.525168 (2023).

39. Y. Liu et al., Preventing oxidative stress: a new role for XBP1. Cell Death Differ 16, 847–857 (2009).

40. S. Yu et al., Glycine ameliorates mitochondrial dysfunction caused by ABT-199 in porcine oocytes. J Anim Sci 99, (2021).

41. Y. Shi et al., Molecular cloning, expression and enzymatic characterization of glutathione S-transferase from Antarctic sea-ice bacteria Pseudoalteromonas sp. ANT506. Microbiol Res 169, 179–184 (2014).

42. M. Tajan et al., Serine synthesis pathway inhibition cooperates with dietary serine and glycine limitation for cancer therapy. Nat Commun 12, 366 (2021).

43. H. Weinstabl et al., Intracellular Trapping of the Selective Phosphoglycerate Dehydrogenase (PHGDH) Inhibitor BI-4924 Disrupts Serine Biosynthesis. J Med Chem 62, 7976–7997 (2019).

44. N. Kiweler et al., Mitochondria preserve an autarkic one-carbon cycle to confer growth-independent cancer cell migration and metastasis. Nat Commun 13, 2699 (2022).

45. R. Hasler et al., Uncoupling of mucosal gene regulation, mRNA splicing and adherent microbiota signatures in inflammatory bowel disease. Gut 66, 2087–2097 (2017).

46. Y. Haberman et al., Ulcerative colitis mucosal transcriptomes reveal mitochondriopathy and personalized mechanisms underlying disease severity and treatment response. Nat Commun 10, 38 (2019).

47. I. Arijs et al., Effect of vedolizumab (anti-alpha4beta7-integrin) therapy on histological healing and mucosal gene expression in patients with UC. Gut 67, 43–52 (2018).

48. S. Saeterstad et al., Profound gene expression changes in the epithelial monolayer of active ulcerative colitis and Crohn’s disease. PLoS One 17, e0265189 (2022).

49. J. Duan et al., Endoplasmic reticulum stress in the intestinal epithelium initiates purine metabolite synthesis and promotes Th17 cell differentiation in the gut. Immunity 56, 1115–1131 e1119 (2023).

50. C. S. Smillie et al., Intra- and Inter-cellular Rewiring of the Human Colon during Ulcerative Colitis. Cell 178, 714–730 e722 (2019).

51. P. M. Irving, S. de Lusignan, D. Tang, M. Nijher, K. Barrett, Risk of common infections in people with inflammatory bowel disease in primary care: a population-based cohort study. BMJ Open Gastroenterol 8, (2021).

52. J. A. Isler, A. H. Skalet, J. C. Alwine, Human cytomegalovirus infection activates and regulates the unfolded protein response. J Virol 79, 6890–6899 (2005).

53. L. Tao et al., Reactive oxygen species oxidize STING and suppress interferon production. Elife 9, (2020).

54. F. V. Marinho, S. Benmerzoug, S. C. Oliveira, B. Ryffel, V. F. J. Quesniaux, The Emerging Roles of STING in Bacterial Infections. Trends Microbiol 25, 906–918 (2017).

55. O. I. Coleman et al., Activated ATF6 Induces Intestinal Dysbiosis and Innate Immune Response to Promote Colorectal Tumorigenesis. Gastroenterology 155, 1539–1552 e1512 (2018).

56. S. Stengel, B. Messner, M. Falk-Paulsen, N. Sommer, P. Rosenstiel, Regulated proteolysis as an element of ER stress and autophagy: Implications for intestinal inflammation. Biochim Biophys Acta Mol Cell Res 1864, 2183–2190 (2017).

57. X. Yin et al., Niche-independent high-purity cultures of Lgr5+ intestinal stem cells and their progeny. Nat Methods 11, 106–112 (2014).

58. J. Dambacher et al., Interleukin 31 mediates MAP kinase and STAT1/3 activation in intestinal epithelial cells and its expression is upregulated in inflammatory bowel disease. Gut 56, 1257–1265 (2007).

59. C. Onyeagocha et al., Latent cytomegalovirus infection exacerbates experimental colitis. Am J Pathol 175, 2034–2042 (2009).

60. K. Hansen et al., Listeria monocytogenes induces IFNbeta expression through an IFI16-, cGAS- and STING-dependent pathway. EMBO J 33, 1654–1666 (2014).

61. L. W. Peterson, D. Artis, Intestinal epithelial cells: regulators of barrier function and immune homeostasis. Nat Rev Immunol 14, 141–153 (2014).

62. R. J. DeBerardinis, N. S. Chandel, Fundamentals of cancer metabolism. Sci Adv 2, e1600200 (2016).

63. A. Kaser, R. S. Blumberg, Endoplasmic reticulum stress and intestinal inflammation. Mucosal Immunol 3, 11–16 (2010).

64. A. Howard et al., Glycine transporter GLYT1 is essential for glycine-mediated protection of human intestinal epithelial cells against oxidative damage. J Physiol 588, 995–1009 (2010).

65. N. Gonen, A. Meller, N. Sabath, R. Shalgi, Amino Acid Biosynthesis Regulation during Endoplasmic Reticulum Stress Is Coupled to Protein Expression Demands. iScience 19, 204–213 (2019).

66. F. F. Diehl, C. A. Lewis, B. P. Fiske, M. G. Vander Heiden, Cellular redox state constrains serine synthesis and nucleotide production to impact cell proliferation. Nat Metab 1, 861–867 (2019).

67. X. Niu et al., Cytosolic and mitochondrial NADPH fluxes are independently regulated. Nat Chem Biol 19, 837–845 (2023).

68. M. Hennequart et al., ALDH1L2 regulation of formate, formyl-methionine, and ROS controls cancer cell migration and metastasis. Cell Rep 42, 112562 (2023).

69. J. Fan et al., Quantitative flux analysis reveals folate-dependent NADPH production. Nature 510, 298–302 (2014).

70. E. Piskounova et al., Oxidative stress inhibits distant metastasis by human melanoma cells. Nature 527, 186–191 (2015).

71. G. S. Ducker et al., Reversal of Cytosolic One-Carbon Flux Compensates for Loss of the Mitochondrial Folate Pathway. Cell Metab 24, 640–641 (2016).

72. P. M. Quiros et al., Multi-omics analysis identifies ATF4 as a key regulator of the mitochondrial stress response in mammals. J Cell Biol 216, 2027–2045 (2017).

73. J. C. van der Mijn et al., Transcriptional and metabolic remodeling in clear cell renal cell carcinoma caused by ATF4 activation and the integrated stress response (ISR). Mol Carcinog 61, 851–864 (2022).

74. D. G. Ryan et al., Disruption of the TCA cycle reveals an ATF4-dependent integration of redox and amino acid metabolism. Elife 10, (2021).

75. J. Han et al., ER-stress-induced transcriptional regulation increases protein synthesis leading to cell death. Nat Cell Biol 15, 481–490 (2013).

76. E. Balsa et al., ER and Nutrient Stress Promote Assembly of Respiratory Chain Supercomplexes through the PERK-eIF2alpha Axis. Mol Cell 74, 877–890 e876 (2019).

77. B. J. Guan et al., Translational control during endoplasmic reticulum stress beyond phosphorylation of the translation initiation factor eIF2alpha. J Biol Chem 289, 12593–12611 (2014).

78. T. M. Seibt, B. Proneth, M. Conrad, Role of GPX4 in ferroptosis and its pharmacological implication. Free Radic Biol Med 133, 144–152 (2019).

79. L. Mayr et al., Dietary lipids fuel GPX4-restricted enteritis resembling Crohn’s disease. Nat Commun 11, 1775 (2020).

80. J. Heijmans et al., ER stress causes rapid loss of intestinal epithelial stemness through activation of the unfolded protein response. Cell Rep 3, 1128–1139 (2013).

81. A. Thavamani, K. K. Umapathi, T. J. Sferra, S. Sankararaman, Cytomegalovirus Infection Is Associated With Adverse Outcomes Among Hospitalized Pediatric Patients With Inflammatory Bowel Disease. Gastroenterology Res 16, 1–8 (2023).

82. E. Mavropoulou et al., Cytomegalovirus colitis in inflammatory bowel disease and after haematopoietic stem cell transplantation: diagnostic accuracy, predictors, risk factors and disease outcome. BMJ Open Gastroenterol 6, e000258 (2019).

83. C. N. Paiva, M. T. Bozza, Are reactive oxygen species always detrimental to pathogens? Antioxid Redox Signal 20, 1000–1037 (2014).

84. F. Ye et al., Reactive oxygen species hydrogen peroxide mediates Kaposi’s sarcoma-associated herpesvirus reactivation from latency. PLoS Pathog 7, e1002054 (2011).

85. M. A. Fisher, M. L. Lloyd, A Review of Murine Cytomegalovirus as a Model for Human Cytomegalovirus Disease-Do Mice Lie? Int J Mol Sci 22, (2020).

86. L. B. Crawford, D. N. Streblow, M. Hakki, J. A. Nelson, P. Caposio, Humanized mouse models of human cytomegalovirus infection. Curr Opin Virol 13, 86–92 (2015).

87. B. B. Madison et al., Cis elements of the villin gene control expression in restricted domains of the vertical (crypt) and horizontal (duodenum, cecum) axes of the intestine. J Biol Chem 277, 33275–33283 (2002).

88. L. Jin et al., MPYS is required for IFN response factor 3 activation and type I IFN production in the response of cultured phagocytes to bacterial second messengers cyclic-di-AMP and cyclic-di-GMP. J Immunol 187, 2595–2601 (2011).

89. A. H. Lee, N. N. Iwakoshi, K. C. Anderson, L. H. Glimcher, Proteasome inhibitors disrupt the unfolded protein response in myeloma cells. Proc Natl Acad Sci U S A 100, 9946–9951 (2003).

90. T. Sato et al., Single Lgr5 stem cells build crypt-villus structures in vitro without a mesenchymal niche. Nature 459, 262–265 (2009).

91. B. Chan et al., The murine cytomegalovirus M35 protein antagonizes type I IFN induction downstream of pattern recognition receptors by targeting NF-kappaB mediated transcription. PLoS Pathog 13, e1006382 (2017).

92. M. T. Sorbara et al., The protein ATG16L1 suppresses inflammatory cytokines induced by the intracellular sensors Nod1 and Nod2 in an autophagy-independent manner. Immunity 39, 858–873 (2013).

93. M. Foroutan et al., Single sample scoring of molecular phenotypes. BMC Bioinformatics 19, 404 (2018).

94. D. D. Bhuva, J. Cursons, M. J. Davis, Stable gene expression for normalisation and single-sample scoring. Nucleic Acids Res 48, e113 (2020).

95. K. Hiller et al., MetaboliteDetector: comprehensive analysis tool for targeted and nontargeted GC/MS based metabolome analysis. Anal Chem 81, 3429–3439 (2009).

